# RABL2 Regulates Ciliation via Controlling IFT-B1 Basal Body Recruitment and ARL3-mediated BBSome Ciliary Retrieval

**DOI:** 10.1101/2022.02.13.480273

**Authors:** Rui-Kai Zhang, Yan-Xia Liu, Wei-Yue Sun, Dong-Xue Bao, Rui-Qin Jia, Chan Zhang, Zhen-Chuan Fan

## Abstract

Highly conserved intraflagellar transport (IFT) trains and certain small GTPases coordinate to direct ciliation and to maintain the ciliary dynamics of signaling molecules via the IFT cargo adaptor BBSome. Unlike murine Rab-like 2 (RABL2) GTPase that enters cilia to drive outward transition zone (TZ) passage of the BBSome, human orthologue fails to enter cilia but resides at the ciliary base, indispensable for ciliation. However, mechanisms underlying how RABL2 regulates ciliation and BBSome barrier passage remain elusive. Here, we show that *Chlamydomonas* RABL2 regulates basal body targeting of the IFT-B1 subcomplex component of IFT trains as a RABL2-specific effector, mediating ciliation via controlling IFT-B1 basal body amount available for assembling anterograde IFT trains. RABL2^GTP^ binds IFT-B1 to perform IFT; sheds from retrograde IFT trains at the proximal ciliary region right above the TZ; and converts to RABL2^GDP^ rapidly. Next, RABL2^GDP^ activates the ciliary membrane anchored Arf-like 3 (ARL3) GTPase (ARL3^GDP^) as a ARL3-specific guanine nucleotide exchange factor. Upon detaching from the ciliary membrane, the active ARL3^GTP^ recruits its BBSome effector, autonomous of retrograde IFT train association, to move cross the TZ for ciliary retrieval. This ensures proper BBSome ciliary turnover for maintaining phototactic response of *Chlamydomonas* cells. For finishing RABL2 ciliary cycle, RABL2^GDP^ passes the TZ for ciliary retrieval by loading onto the ARL3^GTP^/BBSome as a BBSome cargo. Our data thus propose that RABL2 mediates ciliation and BBSome ciliary retrieval simultaneously but via distinct molecular pathways.

**Significance statement:** Intraflagellar transport (IFT) and its cargo adaptor BBSome are indispensable for ciliation and ciliary singling. Rab-like 2 (RABL2) GTPase mediates ciliation and outward transition zone (TZ) passage of BBSomes with mechanisms yet to be determined. Here, we report that RABL2 decides ciliation by controlling the basal body amount of its effector IFT-B1 available for the assembly of anterograde IFT trains. RABL2^GTP^ cycles through cilia as an IFT-B1 cargo; sheds from IFT at the ciliary base; and undergoes nucleotide exchange for activating ARL3 as an ARL3-specfic guanine nucleotide exchange factor. ARL^GTP^ recruits IFT-shed BBSomes to pass the TZ for ciliary retrieval. RABL2^GDP^ exists cilia via ARL3^GTP^/BBSome as a BBSome cargo. Therefore, RABL2 functions both outside and inside cilia for initiating IFT and BBSome ciliary retrieval, respectively.

## Introduction

Cilia (the same as flagella) consist of a microtubule-assembled axoneme surrounded by a specialized ciliary membrane (1, 2). Ciliation relies on intraflagellar transport (IFT) for providing cilia their building precursors and for removing the turnover products out of cilia. During this process, IFT trains, consisting of repeating units of the complexes IFT-A and -B (composed of IFT-B1 and -B2 subcomplexes), traffic along the axoneme in anterograde direction (from the ciliary base to tip) powered by kinesin-2 motor and in retrograde direction (from the ciliary tip to base) driven by cytoplasmic dynein 1b (cytoplasmic dynein 2 in mammals) (3–11). As cellular organelles protruding from eukaryotic cell surface, cilia beat to aid in cell movement or function as antennae to sense extracellular stimuli including odorants, light, mechanical force, and hormones and transduce the signals inside the cell to trigger cellular responses or both. Underlying these signaling is the fact that many G protein-coupled receptors (GPCRs), ion channels and their downstream signaling proteins reside in the ciliary membrane (12–17).

Signaling proteins rely on IFT for entering or/and removing from cilia (12, 15, 18–21). During this process, the BBSome composed of eight BBS proteins (BBS1/2/4/5/7/8/9/18) (7 proteins, BBS1/2/4/5/7/8/9 in *Chlamydomonas reinhardtii*) functions as an adaptor to bridge signaling protein cargoes to IFT-B1 (11, 20, 22–25). BBSomes couple with the anterograde IFT at the basal bodies prior to ciliary entry and anterograde traffic. At the ciliary tip, BBSomes remodel prior to loading onto retrograde IFT trains for trafficking to the ciliary base (19, 26–32). Upon reaching the proximal ciliary region right above the transition zone (TZ), cargo-laden BBSomes shed from retrograde IFT trains and diffuse through the TZ and out of cilia (33). Therefore, defects in BBSome assembly and ciliary cycling cause loss or retention of signaling proteins in the ciliary membrane (22, 23, 34–37). This disrupts ciliary signaling, causing Bardet-Biedl syndrome (BBS), a human ciliopathy characterized by obesity, blindness, kidney failure, and polydactyly (38) and phototactic defects in *C. reinhardtii* (18, 23, 33, 39).

Small GTPases function as molecular switches for controlling BBSome ciliary turnover in various BBSome ciliary cycle steps. For example, *Chlamydomonas* IFT22/BBS3, the heterodimer composed of Rab-like 5 (RABL5) GTPase IFT22 and Arf-like 6 (ARL6) GTPase BBS3, binds the BBSome through a direct interaction between BBS3 and the BBSome (40). IFT22/BBS3 recruits the BBSome to the basal bodies when they both are GTP-bound, controlling BBSome ciliary turnover by deciding its basal body amount available for entering cilia (40). In mammalian cells, BBS3^GTP^ was proposed to enter cilia for GTPase cycling with the aid of RABL4/IFT27 as a BBS3-specific guanine nucleotide exchange factor (GEF) (19). At the ciliary tip, BBS3 reloaded with GTP in turn binds and loads receptor-laden BBSomes onto retrograde IFT trains for trafficking to the ciliary base (19). However, this may not be the case in green algae. *Chlamydomonas* BBS3 promotes cargo (phospholipase D, PLD) association with the BBSome even in a GDP-locked state but does not facilitate BBSome loading onto retrograde IFT trains at the ciliary tip (41). Interestingly, murine RABL2 enters cilia to promote outward TZ passage of GPCR-laden BBSomes for proper ciliary signaling (42). *Chlamydomonas* ARL3 recruits its BBSome effector, autonomous of retrograde IFT train association, to diffuse through the TZ for ciliary retrieval, mediating phototaxis via the BBSome signaling pathway (33). These observations suggest that ARL3 and RABL2 may crosstalk in this biological event.

Besides its role in directing BBSome ciliary removal, RABL2 is indispensable for ciliation in human RPE-hTERT and *Chlamydomonas* cells but not in murine cells (42–45). In RPE-hTERT cells, RABL2 (existing as two high homologous and redundant paralogues RABL2A and RABL2B) binds IFT-B1 as its effector and resides at the basal bodies (43, 44, 46). However, how RABL2 mediates ciliation remains unknown. In this study, we identify RABL2 mediates ciliation via controlling IFT-B1 basal body amount available for assembling anterograde IFT trains in *Chlamydomonas* cells. GTP-loading confers RABL2 to bind IFT-B1 as a cargo for cycling through cilia. Upon retrogradely trafficking to the proximal ciliary region right above the TZ, RABL2^GTP^ sheds from IFT followed by GTPase hydrolysis. RABL2^GDP^ then activates ARL3^GDP^ as a ARL3-specific GEF, mediating phototactic response via promoting outward TZ passage of BBSomes for ciliary retrieval via ARL3^GTP^/BBSome pathway (33). Therefore, RABL2 regulates both ciliation and BBSome ciliary retrieval but via distinct molecular pathways.

## Results

### *Chlamydomonas* RABL2^GTP^ is indispensable for ciliation

*Chlamydomonas* RABL2 shares significant homology with its orthologues in ciliated species and is more closely related to homologues of ciliated non-mammals than mammals and humans phylogenetically (*SI Appendix*, Fig. S1*A* and *B*). Like its human counterpart, *Chlamydomonas* RABL2 is indispensable for ciliation, while RABL2-null murine cells retain cilia of normal length, revealing species- or tissue/cell-specific role of RABL2 for mediating ciliation (42–44, 46). To investigate the effect of *Chlamydomonas* RABL2 nucleotide state on ciliation, we rescued the RABL2-null mutant (LMJ.RY0402.205222) that we named *rabl2^CLiP^* by expressing RABL2 fused at its C-terminus to hemagglutinin (HA) and yellow fluorescent protein (YFP) (RABL2-HA-YFP) and the GTP-locked RABL2^Q80L^-HA-YFP and GDP-locked RABL2^S37N^-HA-YFP variants at the endogenous RABL2 level of parental CC-5325 cells (resulting strains *rabl2 RABL2-HA-YFP*, *rabl2 RABL2^Q83L^-HA-YFP*, and *rabl2 RABL2^S37N^-HA-YFP*) (Fig. 1*A* and *B* and *SI Appendix*, Fig. S1*A*) (44). Unlike the bald (deflagellated) *rabl2^CLiP^, rabl2 RABL2-HA-YFP* and *rabl2 RABL2^Q83L^-HA-YFP* resembled the CC-5325 cell in assembling full-length cilia (Fig. 1*C* and *D*) (44). In contrast, *rabl2 RABL2^S37N^-HA-YFP* remained to be bald (Fig. 1*C* and *D*). To verify these results, we applied vector-based microRNA (miRNA) to knock the endogenous RABL2 down to ~9.0% of wild-type (WT) level in the CC-125 cell; we referred to this strain as RABL2^miRNA^ (Fig. 1*E*). RABL2^miRNA^ assembled cilia ~85% shorter than the CC-125 cell (Fig. 1*F* and *G*). When RABL2 was rescued to WT levels by expressing its N-terminal HA-YFP-tagged versions of various nucleotide states (resulting strains RABL2^Res-WT^, RABL2^Res-Q83L^, and RABL2^Res-S37N^) (Fig. 1*E*), RABL2-HA-YFP and RABL2^Q83L^-HA-YFP rather than RABL2^S37N^-HA-YFP restored cilia to normal length (Fig. 1*F* and *G*). Therefore, GTP-but not GDP-bound RABL2 is indispensable for ciliation in *C. reinhardtii*.

**Figure 1.**
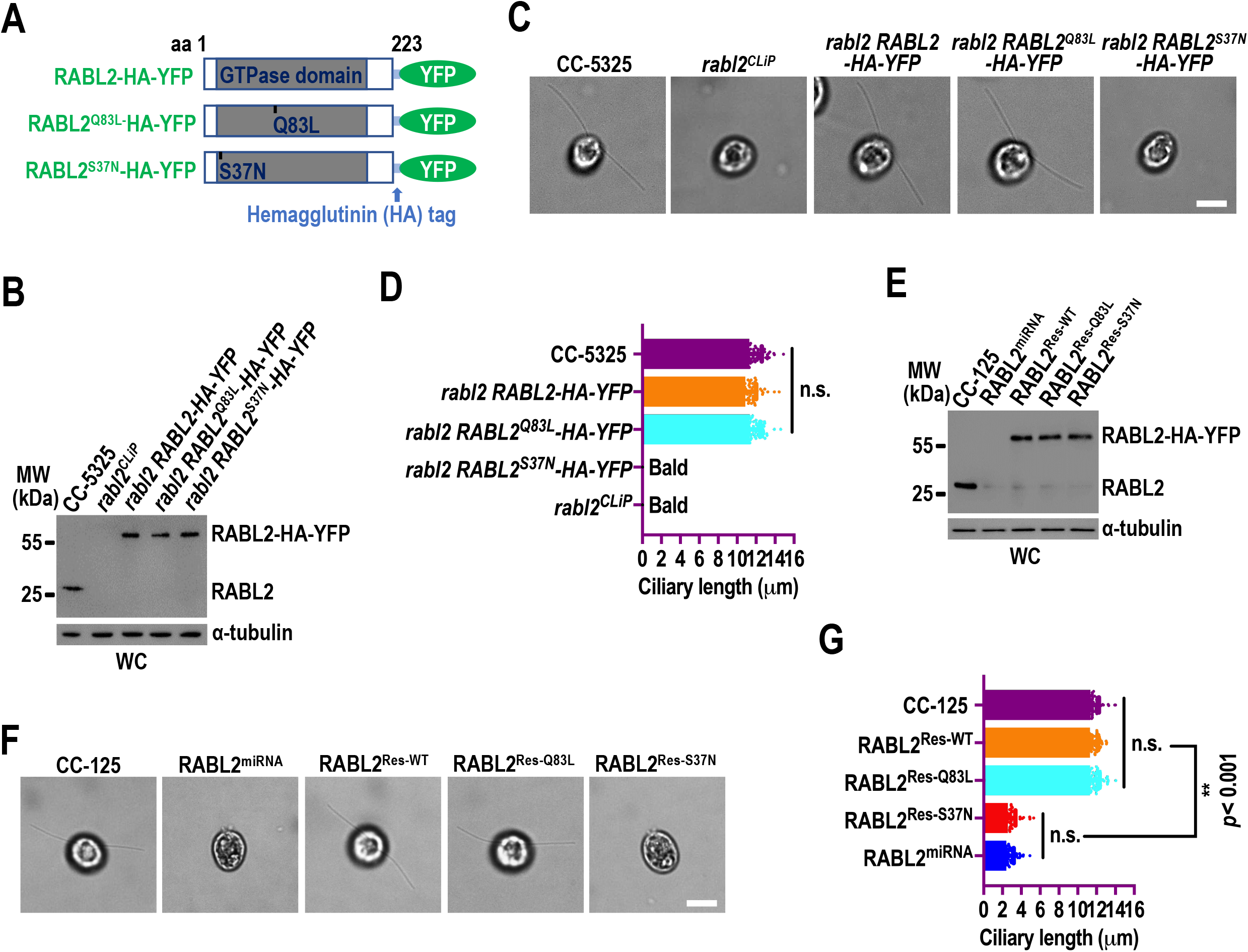
*Chlamydomonas* RABL2^GTP^ is indispensable for ciliation. (*A*) Schematic representation of RABL2-HA-YFP and its GTP- and GDP-locked RABL2^Q83L^-HA-YFP and RABL2^S37N^-HA-YFP variants. aa stands for amino acid. YFP: yellow fluorescence protein. The hemagglutinin (HA) tag was shown. (*B*) Immunoblots of whole cell (WC) samples of cells indicated on the top probed with α-RABL2. (*C*) Representative phase contrast (PC) images of cells in panel *B* showing cilia. Scale bars: 5 μm. (*D*) *rabl2 RABL2-HA-YFP* (10.88 ± 1.15 μm, n = 100) and *rabl2 RABL2^Q83L^-HA-YFP* (11.31 ± 1.14 μm, n = 100) cells had full length cilia as compared to CC-5325 cells (11.28 ± 1.31 μm, n = 100). *rabl2^CLiP^* and *rabl2 RABL2^S37N^-HA-YFP* cells were bald. (*E*) Immunoblots of WC samples of cells indicated on the top probed with α-RABL2. (*F*) Representative PC images of cells shown in panel *E* showing cilia. Scale bars: 10 μm. (*G*) RABL2^Res-WT^ (11.29 ± 0.90 μm, n = 100) and RABL2^Res-Q83L^ (11.31 ± 1.05 μm, n = 100) cells had full length cilia as compared to CC-125 cells (11.35 ± 0.91 μm, n = 100). RABL2^miRNA^ (2.54 ± 0.79 μm, n = 100) and RABL2^Res-S37N^ (1.99 ± 0.80 μm, n = 100) cells had cilia with similar length which both are much shorter than the WT cells. For panels *B* and *E*, α-tubulin was used for adjusting the loading. MW stands for molecular weight. For panels *D* and *G*, mean lengths are listed. Error bar indicates S.D. and “n” indicates the number of cilia counted. n.s.: non-significance. **: significance at *p* < 0.001.

### RABL2 controls IFT-B1 basal body recruitment

RABL2 interacts with its IFT-B1 effector via binding the heterodimeric IFT74/81 component of IFT-B1 directly in a GTP-dependent manner and resides at the ciliary base in human and murine cells in the steady state (42–46). We wondered if RABL2^GTP^ binds IFT-B1 in cytoplasm and recruits it to the basal bodies dynamically. To test this notion, immunoprecipitation assays were performed on cell body samples of *rabl2 RABL2-HA-YFP*, *rabl2 RABL2^Q83L^-HA-YFP*, and *rabl2 RABL2^S37N^-HA-YFP* with the HA-YFP-expressing HR-YFP cell being used as a negative control (47). It has been shown that IFT-A, IFT-B1, and IFT-B2 subcomplexes components of IFT trains become separated from one another in HMEKN buffer (Fig. 2*A*) (39). In the same buffer, RABL2^Q83L^-HA-YFP immunoprecipitated IFT-B1 (represented by IFT46 and IFT70) but not IFT-A (represented by IFT43 and IFT139) and IFT-B2 (represented by IFT38 and IFT57) (Fig. 2*B*). RABL2-HA-YFP resembled RABL2^S37N^-HA-YFP for recovering none of them, consistent with the human and murine studies showing that RABL2^GDP^ exists in cytoplasm but converts to RABL2^GTP^ for IFT-B1 association (Fig. 2*B*) (42–44). Supportive of this notion, the majority of RABL2^Q83L^-HA-YFP co-sedimented with IFT-B1 in sucrose density gradient of the cell body and RABL2-HA-YFP and RABL2^S37N^-HA-YFP did not do so (Fig. 2*C*). This notion was further confirmed as the endogenous RABL2 co-fractionated with IFT-B1 in the presence of the nonhydrolyzable GTP analog GTPγS that locks RABL2 in a GTP-bound state but did not do so in the absence of nucleotides or the presence of GDP that locks RABL2 in a GDP-bound state (Fig. 2*D*). Since IFT74/81 can pull down RABL2 only in the presence of GTPγS and RBAL2^Q83L^ but not RABL2^S37N^ *in vitro*, we conclude that *Chlamydomonas* RABL2 interacts with IFT-B1 via directly binding IFT74/81 in a GTP-dependent manner, consistent with the human and murine data (Fig. 2*E*). (42, 43).

**Figure 2.**
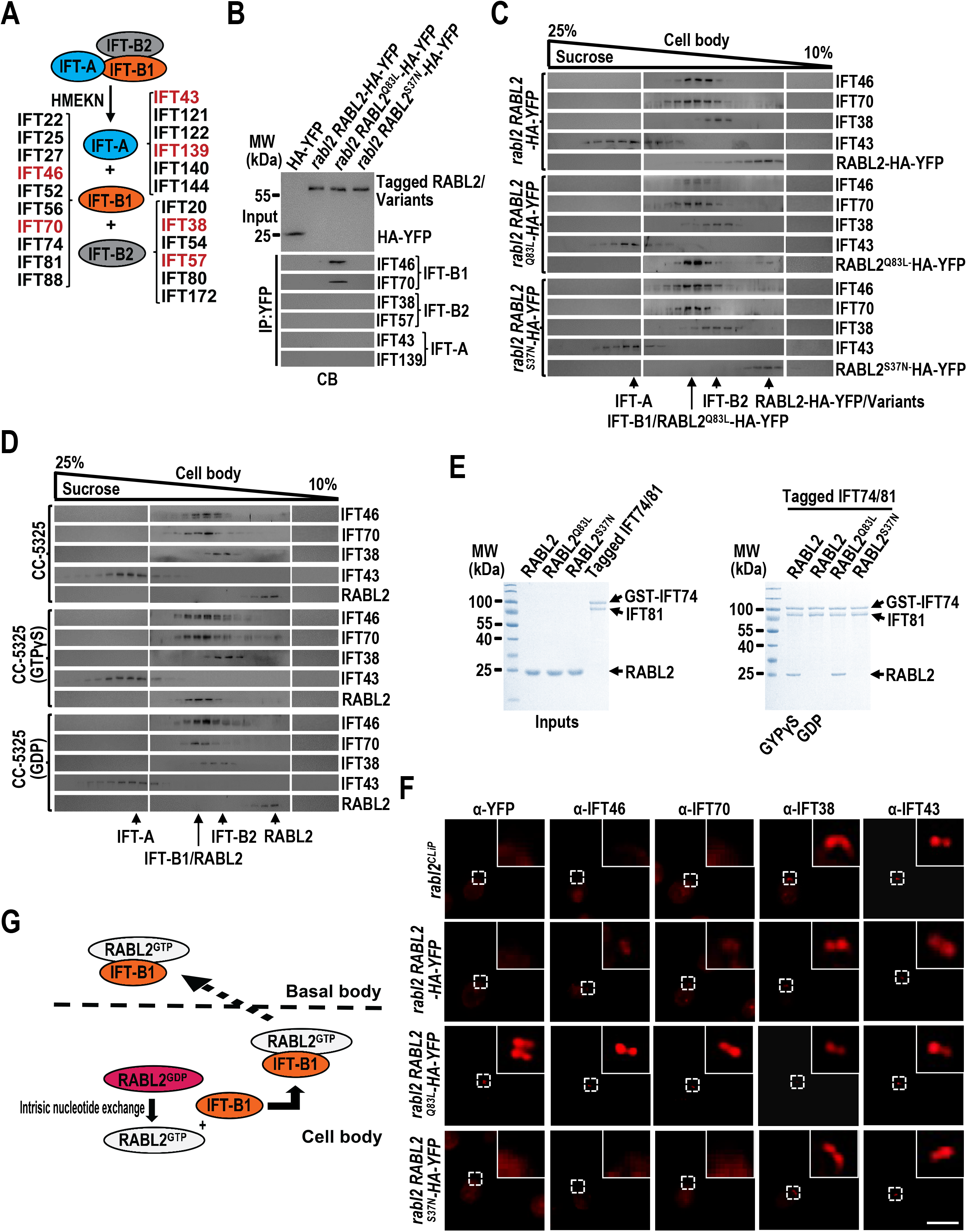
RABL2 controls IFT-B1 basal body recruitment. (*A*) Schematic representation showing that IFT subcomplexes IFT-A, IFT-B1, and IFT-B2 remain to be separated from one another in HMEKN buffer. The IFT subcomplex subunits were shown. The IFT subunits in red were probed as shown in penal *B*. (*B*) Immunoblots of α-YFP-captured proteins from cell body (CB) samples (in HMEKN buffer) of cells indicated on the top probed for the IFT-A subunits IFT43 and IFT139, the IFT-B1 subunits IFT46 and IFT70, and the IFT-B2 subunits IFT38 and IFT57. The input was adjusted with α-YFP by immunoblotting. (*C*) Immunoblots of sucrose density gradient of CB samples of cells indicated on the left probed with α-IFT46, α-IFT70, α-IFT38, α-IFT43, and α-YFP. (*D*) Immunoblots of sucrose density gradient of CB samples of CC-5325 in the presence of GTPγS, GDP, or none of them probed with α-IFT46, α-IFT70, α-IFT38, α-IFT43, and α-RABL2. (*E*) Bacterially expressed RABL2, RABL2^Q83L^, RABL2^S37N^, and GST-IFT74/81 purified before SDS-PAGE electrophoresis and Coomassie staining (left panel). GST-IFT74/81 mixed with RABL2 (in the presence of GTPγS or GDP), RABL2^Q83L^, and RABL2^S37N^, complexes recovered on glutathione beads before SDS-PAGE electrophoresis and Coomassie staining (right panel). (*F*) Cells indicated on the left stained with α-YFP, α-IFT46, α-IFT70, α-IFT38, and α-IFT43. Inset shows the basal body and the transition zone staining. Scale bars: 10 μm. (*G*) Schematic representation of how RABL2 binds and recruits IFT-B1 as its effector to the basal body. For panels *B* and *E*, MW stands for molecular weight.

As performed on *rabl2 RABL2-HA-YFP*, *rabl2 RABL2^Q83L^-HA-YFP*, and *rabl2 RABL2^S37N^-HA-YFP*, immunostaining failed to visualize RABL2-HA-YFP and RABL2^S37N^-HA-YFP at the basal bodies (Fig. 2*F*). RABL2^Q83L^-HA-YFP, however, was easily detected to enrich at the basal bodies, revealing that GTP-loading confers more RABL2 to traffic to the basal bodies (Fig. 2*F*). As compared to RABL2-HA-YFP and RABL2^S37N^-HA-YFP, RABL2^Q83L^-HA-YFP was also enriched at the ciliary base obviously above the basal bodies, recapitulating rodent RABL2^GTP^ in accumulating at the ciliary base right above the TZ (Fig. 2*F*) (42). Furthermore, IFT-A (represented by IFT43) and IFT-B2 (represented by IFT38) both resided at the basal bodies of all three strains at *rabl2^CLiP^* levels, while IFT-B1 (represented by IFT46 and IFT70) targeted to the basal bodes of *rabl2 RABL2-HA-YFP* and *rabl2 RABL2^Q83L^-HA-YFP* but not *rabl2 RABL2^S37N^-HA-YFP* and *rabl2^CLiP^* (Fig. 2*F*). Even for the latter cells, IFT-B1 resided at the basal bodies in *rabl2 RABL2^Q83L^-HA-YFP* much more than in *rabl2 RABL2-HA-YFP* (Fig. 2*F*). Based on these evidence, we conclude that *Chlamydomonas* RABL2 resembles its human orthologue to exist in a GDP-bound state (RABL2^GDP^) in cytoplasm (Fig. 2*G*) (43). Upon activated by intrinsic nucleotide exchange (43), RABL2^GDP^ converts to RABL2^GTP^, which recruits its IFT-B1 effector to the basal bodies (Fig. 2*G*).

### RABL2^GTP^ can shed from retrograde IFT trains but fails to pass the TZ for ciliary retrieval

Human RABL2^GTP^ initiates IFT at the ciliary base followed by dissociation with IFT-B1 by GTP hydrolysis, thus failing to enter cilia itself (43, 44, 46). In contrast, murine RABL2^GTP^ but not RABL2^GDP^ undergoes IFT as an IFT-B1 cargo, sheds from retrograde IFT trains at the ciliary base, and fails to pass the TZ for ciliary retrieval (42). *Chlamydomonas* RABL2^GTP^ resembles its murine counterpart in accumulating at the ciliary base, while how it behaves like this way in cilia remains unknown (Fig. 2*F*). Reflecting its buildup at the ciliary base (Fig. 2*F*), RABL2^Q83L^-HA-YFP was determined to retain in cilia ~6.5-fold higher than RABL2-HA-YFP and the endogenous RABL2 in the steady state (Fig. 3*A*). RABL2-HA-YFP and RABL^Q83L^-HA-YFP resembled the endogenous RABL2 in residing in the ciliary matrix where IFT proteins reside, signifying RABL2 interaction with IFT (*SI Appendix*, Fig. S2*A*). Reflecting RABL2 low ciliary abundance relative to RABL2^Q83L^-HA-YFP (Fig. 3*A*), RABL2-HA-YFP was undetectable by total internal reflection inverted fluorescence (TIRF) even when its YFP tag was replaced with as many as three copies of mNeonGreen (3NG) (RABL2-HA-3NG), which is supposed to be more brighter than a single-copy of YFP (Fig. 3*B* and *SI Appendix*, Fig. S2*B* and *C*) (42). RABL2^Q83L^-HA-YFP, however, was easily visualized to undergo IFT at bidirectional frequencies both at ~70% of IFT trains represented by IFT46-YFP of *ift46-1 IFT46-YFP* cells (Fig. 3*B-D* and Movies S1 and S2) (48). This excluded RABL2^GTP^ ciliary accumulation from being caused by its unaltered IFT behavior. Accordingly, RABL2^Q83L^-HA-YFP buildup in cilia did not reflect its increased basal body amount but could be an outcome of its inability to pass the TZ for ciliary retrieval after shedding from the retrograde IFT trains (Fig. 2*F*) (42). This notion was verified as RABL2^Q83L^-HA-YFP was indeed found to enrich at the CEP290-labeled TZ (Fig. 3*E*).

**Figure 3.**
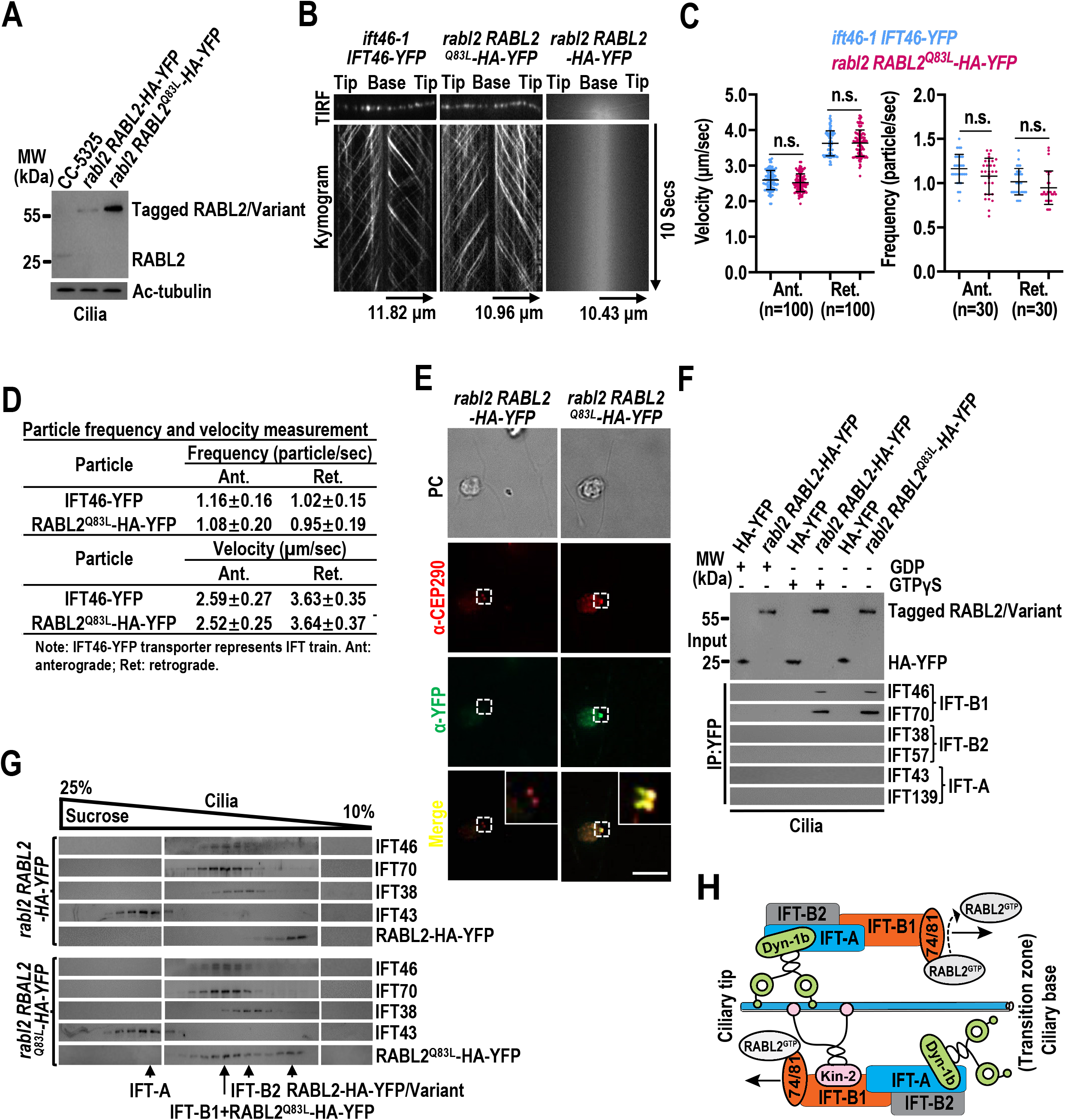
RABL2^GTP^ performs normal IFT but fails to pass the TZ for ciliary retrieval. (*A*) Immunoblots of cilia of cells indicated on the top probed with α-RABL2. Acetylated (Ac)-tubulin was used as a loading control. (*B*) TIRF images and corresponding kymograms of cells indicated on the top (Movies S1-S3, 15 fps). The time and transport lengths are indicated on the right and on the bottom, respectively. The ciliary base (base) and tip (tip) were shown. (*C* and *D*) Velocities and frequencies of fluorescence proteins to traffic inside cilia of cells in panel *B* shown as graphs (*C*) and numbers (*D*). Mean values are listed. Error bar indicates S.D. n: number of cilia analyzed. n.s.: non-significance. (*E*) Cells indicated on the top stained with α-CEP290 (red) and α-YFP (green). Inset shows the basal body and the transition zone staining. PC images of cells were shown. Scale bars: 10 μm. (*F*) Immunoblots of α-YFP-captured proteins from HA-YFP and *rabl2 RABL2-HA-YFP* cilia in the presence of GDP or GTPγS and *rabl2* RABL2^Q83L^-HA-YFP cilia in the absence of GDP and GTPγS probed with α-IFT46, α-IFT70, α-IFT38, α-IFT57, α-IFT43, and α-IFT139. The input was adjusted with α-YFP by immunoblotting. (*G*) Immunoblots of sucrose density gradient of cilia of cells indicated on the left probed with α-IFT46, α-IFT70, α-IFT38, α-IFT43, and α-YFP. (*H*) Schematic representation of how RABL2^GTP^ couples with IFT-B1 through directing binding IFT74/81 for performing IFT and sheds from retrograde IFT at the ciliary base right above the TZ. For panels *A* and *F*, MW stands for molecular weight.

To dissect the molecular base of how RABL2 binds IFT in cilia, we performed immunoprecipitation on *rabl2 RABL2-HA-YFP* and *rabl2 RABL2^Q83L^-HA-YFP* cilia. RABL2^Q83L^-HA-YFP recovered IFT-B1 alone, consistent with the observation that RABL2-HA-YFP immunoprecipitated IFT-B1 but not IFT-A and IFT-B2 in the presence of GTPγS (Fig. 3*F*). Accordingly, in the presence of GTPγS, the endogenous RABL2 co-fractionated with IFT-B1 alone in CC-5325 cilia, consistent with the observation that RABL2^Q83L^-HA-YFP co-sedimented with IFT-B1 but not IFT-A and IFT-B2 in *rabl2 RABL2^Q83L^-HA-YFP* cilia (Fig. 3*G* and *SI Appendix*, Fig. S2*D*). In this case, partial RABL2^Q83L^-HA-YFP remained to be separated from IFT-B1, representing the RABL2^GTP^ portion that sheds from retrograde IFT trains but fails to pass the TZ for ciliary retrieval (Fig. 3*G*) (42). In summary, RABL2^GTP^ undergoes IFT by acting as an IFT-B1 cargo (Fig. 3*H*). RABL2^GTP^ can shed from the IFT-B1 subcomplex component of retrograde IFT trains right above the TZ but fails to pass the diffusion barrier at the TZ for ciliary retrieval, consistent with the murine study (Fig. 3*H*) (42). Remarkedly, RABL2-HA-YFP resembled the endogenous RABL2 in keeping separated from IFT-A, IFT-B1, and IFT-B2 in cilia, indicating that RABL2^GTP^, once shedding from retrograde IFT trains at the ciliary base right above the TZ, can convert to RABL2^GDP^ rapidly with a mechanism yet to be determined (Fig. 3*G* and *SI Appendix*, Fig. S2*D*).

### RABL2 and the BBSome interdepend on one another for outward TZ passage

Murine RABL2^D73G^ variant causes infertility and obesity in mice but does not affect ciliation (42, 45, 49). This variant can recruit IFT-B1 to the basal bodies for IFT initiation and ciliation but fails to enter cilia itself (42). We accordingly generated a RABL2^D76G^-HA-YFP-expressing *rabl2^CLiP^* strain that we named *rabl2 RABL2^D76G^-HA-YFP* (*SI Appendix*, Fig. S1*A* and S3*A*). When expressed at RABL2 level of parental CC-5325 cells, RABL2^Q76G^-HA-YFP resembled RABL2-HA-YFP in restoring ciliation, while itself as expected failed to enter cilia (Fig. 1*C*, 4*A*, and *SI Appendix*, Fig. S3*B* and *C*). As reflected by normal ciliation, *rabl2 RABL2^D76G^-HA-YFP* resembled *rabl2 RABL2-HA-YFP* in retaining IFT proteins at normal levels in both whole cell samples and cilia (Fig. 4*A* and *SI Appendix*, Fig. S3*B* and *C*). Like RABL2^Q83L^-HA-YFP, RABL2^D76G^-HA-YFP was bound to IFT-B1 rather than IFT-A and IFT-B2 in a GTP-dependent manner in cytoplasm (Fig. 2*B*, *SI Appendix*, Fig. S3*D*). The data suggest that RABL2^D76G^ can recruit IFT-B1 to the basal bodies for initiating normal IFT for ciliation but becomes defective in ciliary entry.

**Figure 4.**
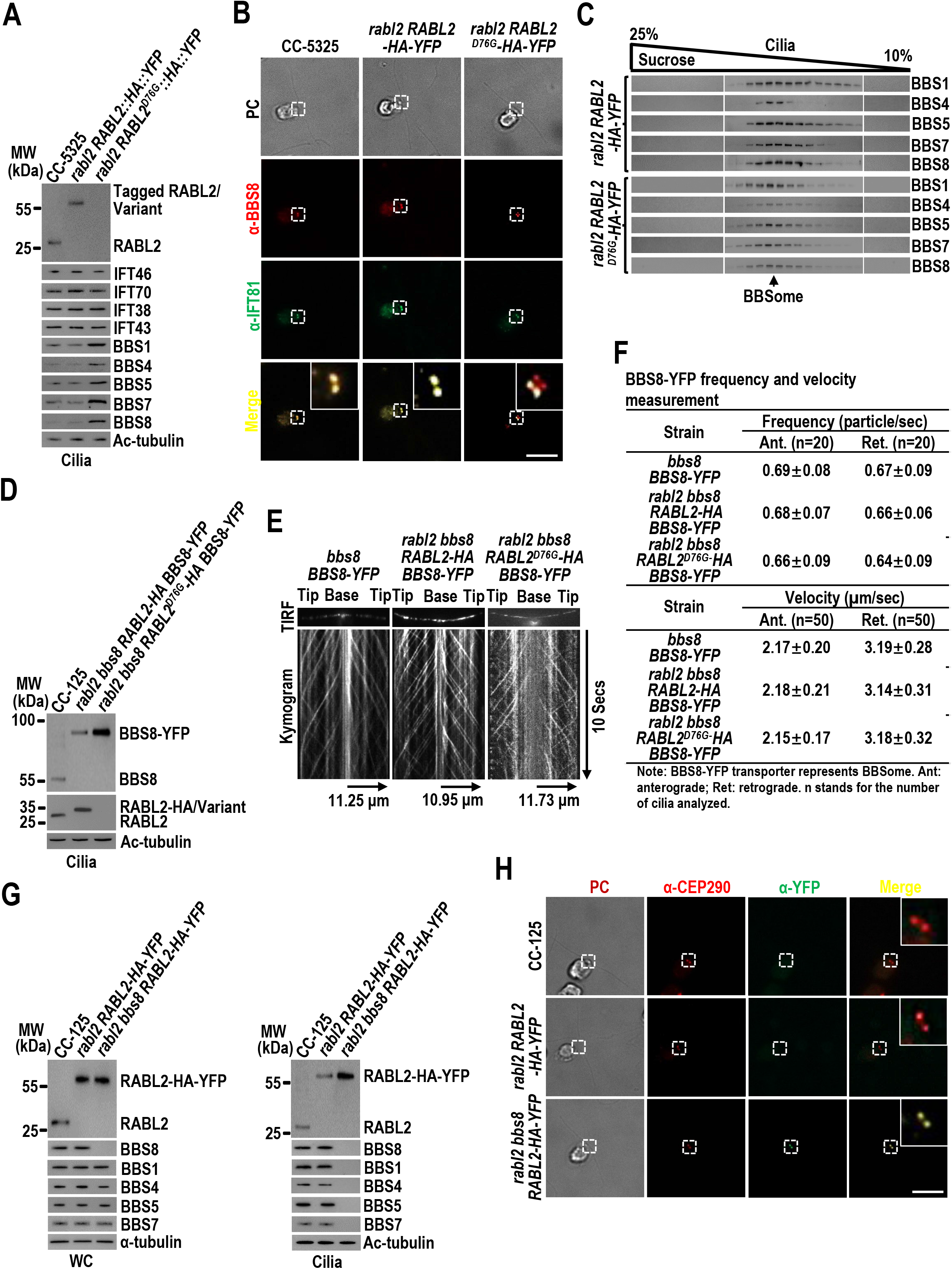
RABL2 and the BBSome interdepend on one another for outward TZ passage. (*A*) Immunoblots of cilia of cell indicated on the top probed with α-RABL2, α-IFT46, α-IFT70, α-IFT38, α-IFT43, α-BBS1,α-BBS4, α-BBS5, α-BBS7, and α-BBS8. (*B*) Cells indicated on the top stained with α-BBS8 (red) and α-IFT81 (green). (*C*) Immunoblots of sucrose density gradient of cilia of cells indicated on the left probed for BBS1, BBS4, BBS5, BBS7, BBS8. (*D*) Immunoblots of cilia of cells indicated on the top probed with α-BBS8 and α-RABL2. (*E*) TIRF images and corresponding kymograms of cells indicated on the top (Movies S6-S8, 15 fps). The time and transport lengths are indicated on the right and on the bottom, respectively. The ciliary base (base) and tip (tip) were shown. (*F*) Velocities and frequencies of fluorescence proteins to traffic inside cilia of cells in panel *E* shown as numbers. Mean values are listed. Error bar indicates S.D. from 100 and 30 cilia analyzed for determining velocity and frequency, respectively. (*G*) Immunoblots of WC samples and cilia of cells indicated on the top probed with α-RABL2, α-BBS8, α-BBS1, α-BBS4, α-BBS5, and α-BBS7. (*H*) Cells indicated on the left stained with α-BBS8 (red) and α-YFP (green). For panels *A*, *D*, and *G*, α-tubulin and acetylated (Ac)-tubulin were used as a loading control for WC samples and cilia, respectively. MW stands for molecular weight. For panels *B* and *H*, inset shows the basal body and the transition zone staining. PC images of cells were shown. Scale bars: 10 μm.

Murine RABL2 promotes outward TZ passage of BBSomes for ciliary retrieval (42). To investigate if this applies in *Chlamydomonas* cells, we examined *rabl2 RABL2-HA-YFP* and *rabl2 RABL2^D76G^-HA-YFP*. As revealed by immunoblotting, they both retained the BBSome (represented by BBS1, BBS4, BBS5, BBS7, and BBS8) at normal level (*SI Appendix*, Fig. S3*B*). However, *rabl2 RABL2^D76G^-HA-YFP* accumulated the BBSome in cilia to a level ~3.5-fold higher than *rabl2 RABL2-HA-YFP* and CC-5325 (Fig. 4*A*) (42). Replacement of RABL2 with RABL2^D76G^-HA-YFP did not influence BBSome basal body targeting nor its integrity in cytoplasm and cilia (Fig. 4*B* and *C*, *SI Appendix*, Fig. S3*E*). It just conferred the BBSome (represented by BBS8) to build up at the ciliary base above the IFT81 and BBS8 double labeled basal bodies (Fig. 4*B*). To discern if this BBSome buildup was caused by its altered IFT behavior, we generated strains *rabl2 bbs8 RABL2-HA BBS8-YFP* and *rabl2 bbs8 RABL2^D76G^-HA BBS8-YFP* (*SI Appendix*, Fig. S3*F*) (33). Two strains had BBS8 replaced with BBS8-YFP (*SI Appendix*, Fig. S3*F*). As expected, replacement of RABL2 with RABL2^D76G^-HA caused BBSome buildup in cilia (Fig. 4*D* and *SI Appendix*, Fig. S3*F*). As assayed by TIRF, the BBSome (represented by BBS8-YFP) performed IFT at bidirectional frequencies in *rabl2 bbs8 RABL2-HA BBS8-YFP* and *rabl2 bbs8 RABL2^D76G^-HA BBS8-YFP* the same as in *bbs8 BBS8-YFP*, excluding the loss of RABL2 in cilia from altering the IFT behavior of the BBSome (Fig. 4*E* and *F*, *SI Appendix*, Fig. S3*G*, and Movies S3-S5). Therefore, RABL2 does not affect BBSome shedding from retrograde IFT trains at the ciliary base but is required for it to pass the TZ for ciliary retrieval (33).

To discern if RABL2 requires the BBSome for ciliary retrieval, we generated a RABL2-HA-YFP-expressing *rabl2-* and *bbs8-* double null strain *rabl2 bbs8 RABL2-HA-YFP*, which expresses RABL2-HA-YFP at the endogenous RABL2 level of CC-5325 cells (Fig. 4*G*). Knockout of BBS8 causes BBSome disassembly in cytoplasm so that this strain was deprived of the BBSome in cilia (Fig. 4*G*) (33). Of note, RABL2-HA-YFP accumulated in *rabl2 bbs8 RABL2-HA-YFP* cilia to a level ~6.5-fold higher than in *rabl2 RABL2-HA-YFP* cilia and its buildup was defined to the CEP290-labeled TZ (Fig. 4*H*). Since the loss of the BBSome in cilia does not affect IFT (23), we conclude that, upon shedding from retrograde IFT trains at the ciliary base, RABL2 requires the BBSome for passing the TZ and out of cilia.

### RABL2^GDP^ promotes outward TZ passage of the BBSome via activating ARL3

ARL3 recruits the BBSome, autonomous of retrograde IFT train association, as its effector to diffuse through the TZ and out of cilia (33). Accordingly, the loss of ARL3 causes BBSome buildup at the ciliary base (33). Since the loss of RABL2 in cilia leads to the same BBSome retention pattern, we reasonably asked if RABL2 mediates outward TZ passage of the BBSome via ARL3. To answer this question, we examined the ARL3-null *arl3^CLiP^* cell and the three rescuing strains *arl3 ARL3-HA-YFP, arl3 ARL3^Q70L^-HA-YFP*, and *arl3 ARL3^T30N^-HA-YFP*, which express ARL3-HA-YFP in a WT, GTP-, and GDP-locked state, respectively (33). ARL3 knockout did not alter cellular levels of the BBSome and RABL2, while they both accumulated in *arl3 ARL3^T30N^-HA-YFP* cilia but not *arl3 ARL3-HA-YFP* and *arl3 ARL3^Q70L^-HA-YFP* cilia (Fig. 5*A* and *SI Appendix*, Fig. S4*A*). This is easy to understand as the BBSome relies on ARL3^GTP^ for passing the TZ for ciliary retrieval and is required for RABL2 to pass the TZ for ciliary retrieval (Fig. 4 and *SI Appendix*, Fig. S4*B*) (33).

**Figure 5.**
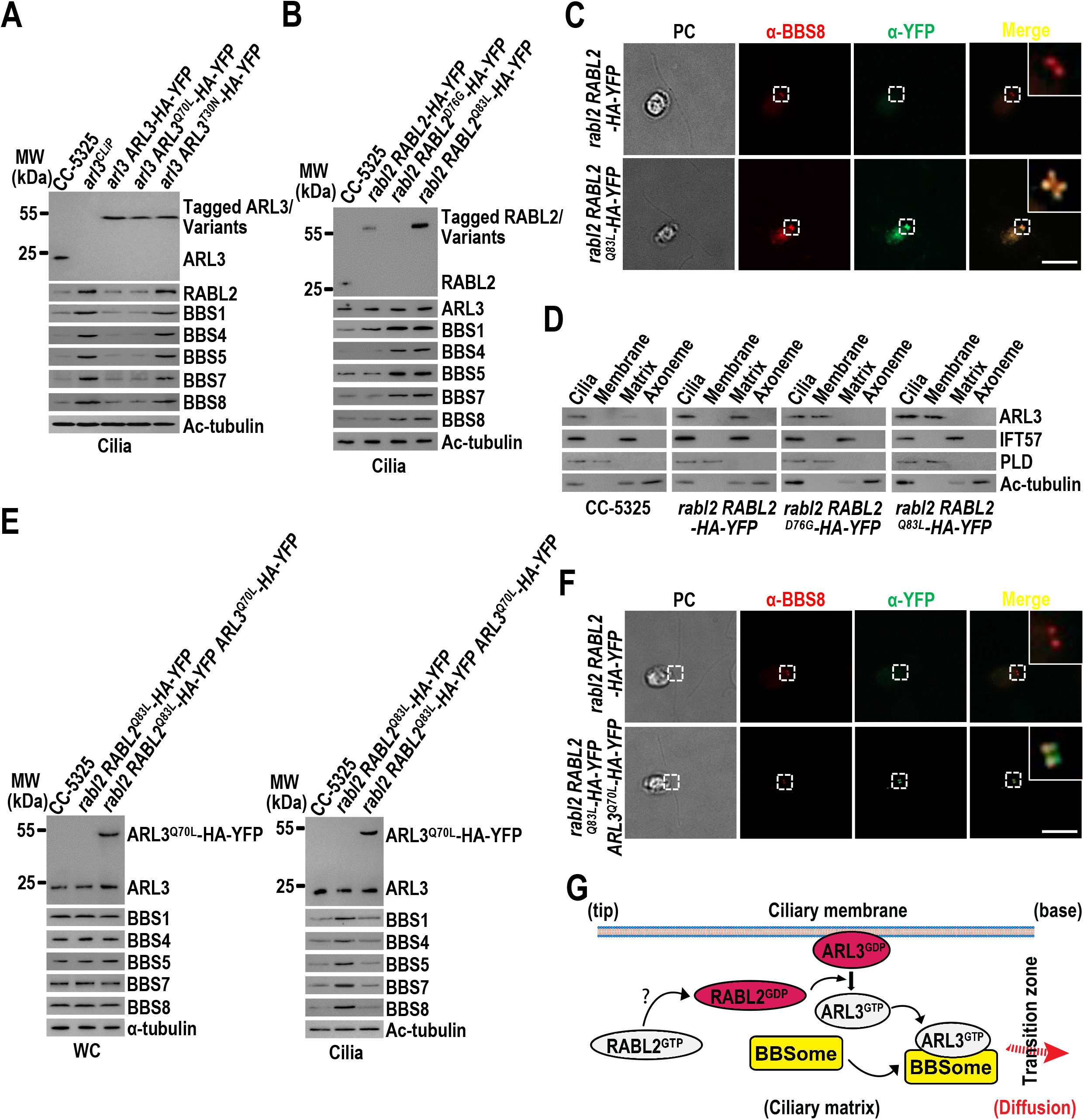
RABL2^GDP^ promotes outward TZ passage of the BBSome via activating ARL3. (*A*) Immunoblots of cilia of cells indicated on the top probed with α-ARL3, α-RABL2, α-BBS1, α-BBS4, α-BBS5, α-BBS7, and α-BBS8. (*B*) Immunoblots of cilia of cells indicated on the top probed with α-RABL2, α-ARL3, α-BBS1, α-BBS4, α-BBS5, α-BBS7, and α-BBS8. (*C*) Cells indicated on the left stained with α-BBS8 (red) and α-YFP (green). (*D*) Immunoblots of ciliary fractions of cells indicated on the bottom probed with α-ARL3, α-IFT57 (ciliary matrix marker), α-PLD (ciliary membrane marker), and acetylated (Ac)-tubulin (axoneme marker). (*E*) Immunoblots of WC samples and cilia of cells indicated on the top probed with α-ARL3, α-BBS1, α-BBS4, α-BBS5, α-BBS7, and α-BBS8. (*F*) Cells indicated on the left stained with α-BBS8 (red) and α-YFP (green). (*G*) Schematic representation of how RABL2 activates ARL3 for promoting outward TZ passage of the BBSome. ? stands for an unknown mechanism. For panels *A*, *B*, and *E*, α-tubulin and acetylated (Ac)-tubulin were used as a loading control for WC samples and cilia, respectively. MW stands for molecular weight. For panels *C* and *F*, inset shows the basal body and the transition zone staining. PC images of cells were shown. Scale bars: 10 μm.

*rabl2 RABL2-HA-YFP, rabl2 RABL2^D76G^-HA-YFP*, and *rabl2 RABL2^Q83L^-HA-YFP* all retained ARL3 at normal levels in both whole cell samples and cilia, revealing that ARL3 does not require the BBSome for maintaining its ciliary turnover (Fig. 5*B* and *SI Appendix*, Fig. S4*C*) (33). We noticed that the loss of RABL2 in cilia as shown by *rabl2 RABL2^D76G^-HA-YFP* caused the BBSome to accumulate at the ciliary base (Fig. 4*A* and *B*). RABL2-HA-YFP restored BBSome ciliary abundance (represented by BBS8) back to normal (Fig. 5*B* and *C*). Replacement of RABL2 with RABL2^Q83L^-HA-YFP, however, failed to do so (Fig. 5*B* and *C*). We have known previously that ARL3^GDP^, upon diffusing into cilia by binding the ciliary membrane, rapidly converts into ARL3^GTP^ for detaching from the ciliary membrane so as to reside in the ciliary matrix (33). Since ARL3 resided in the ciliary matrix of CC-5325 and *rabl2 RABL2-HA-YFP* but remained to bind the ciliary membrane of *rabl2 RABL2^D76G^-HA-YFP* and *rabl2 RABL2^Q83L^-HA-YFP*, we conclude that the loss of RABL2 in cilia and the replacement of RABL2 with RABL2^Q83L^ both can prevent ARL3^GDP^ from converting to ARL3^GTP^ in cilia (Fig. 5*D*). As reflected by these results, RABL2 is essential for ARL3 activation in cilia, while RABL2^GTP^ fails to do so. When ARL3^Q70L^-HA-YFP was expressed in *rabl2 RABL2^Q83L^-HA-YFP* (resulting strain *rabl2 RABL2^Q83L^-HA-YFP ARL3^Q70L^-HA-YFP*), it competed with the endogenous ARL3 for entering cilia and can almost fully restore the BBSome accumulation at the TZ region back to normal (Fig. 5*E* and *F*). Taking together the result that RABL2^GTP^ rapidly converts to RABL2^GDP^ after shedding from retrograde IFT trains at the ciliary base (Fig. 3*G* and *SI Appendix*, Fig. S2*D*), we conclude that it is RABL2^GDP^ that promotes the outward TZ passage of the BBSome for ciliary retrieval by activating ARL3 (Fig. 5*G*) (33).

### RABL2 is a ARL3-specific GEF

RABL2 catalyzes ARL3 for converting from being GDP-bound to being GTP-bound, identifying RABL2 could be a ARL3-specific GEF. To explore if RABL2 interacts with ARL3, we performed immunoprecipitation on cilia of *arl3 bbs8 ARL3-HA-YFP, arl3 bbs8 ARL3^Q70L^-HA-YFP*, and *arl3 bbs8 ARL3^T30N^-HA-YFP* with the HA-YFP-expressing HR-YFP cell being used as a negative control (47). Three cells did not contain the BBSome in cilia owing to the lack of intact BBSome entities in cytoplasm caused by BBS8 knockout (33). In the presence of GTPγS that locks RABL2 and ARL3 both in a GTP-bound state, ARL3-HA-YFP and its variants all failed to recover RABL2 (Fig. 6*A*). ARL3^T30N^-HA-YFP but not ARL3^Q70L^-HA-YFP resembled ARL3-HA-YFP in recovering RABL2 in the presence of GDP that locks two GTPases in a GDP-bound state simultaneously (Fig. 6*A*). We next generated *rabl2 bbs8 RABL2^Q83L^-HA-YFP*, which expresses RABL2^Q83L^-HA-YFP at the WT RABL2 level of CC-125 cells at the RABL2- and BBS8 double-null *rabl2 bbs8* genetic background (Fig. 6*B*). In the absence of the BBSome, RABL2-HA-YFP and RABL2^Q83L^-HA-YFP accumulated in cilia as expected (Fig. 6*B*). They both failed to immunoprecipitate ARL3 in cilia in the presence of GTPγS (Fig. 6*C*). The GDP addition conferred RABL2-HA-YFP but not RABL2^Q83L^-HA-YFP to recover ARL3 (Fig. 6*C*). These findings suggest that RABL2 interacts with ARL3 in cilia only when they both are being GDP-bound. They interact with one another in a BBSome-independent manner.

**Figure 6.**
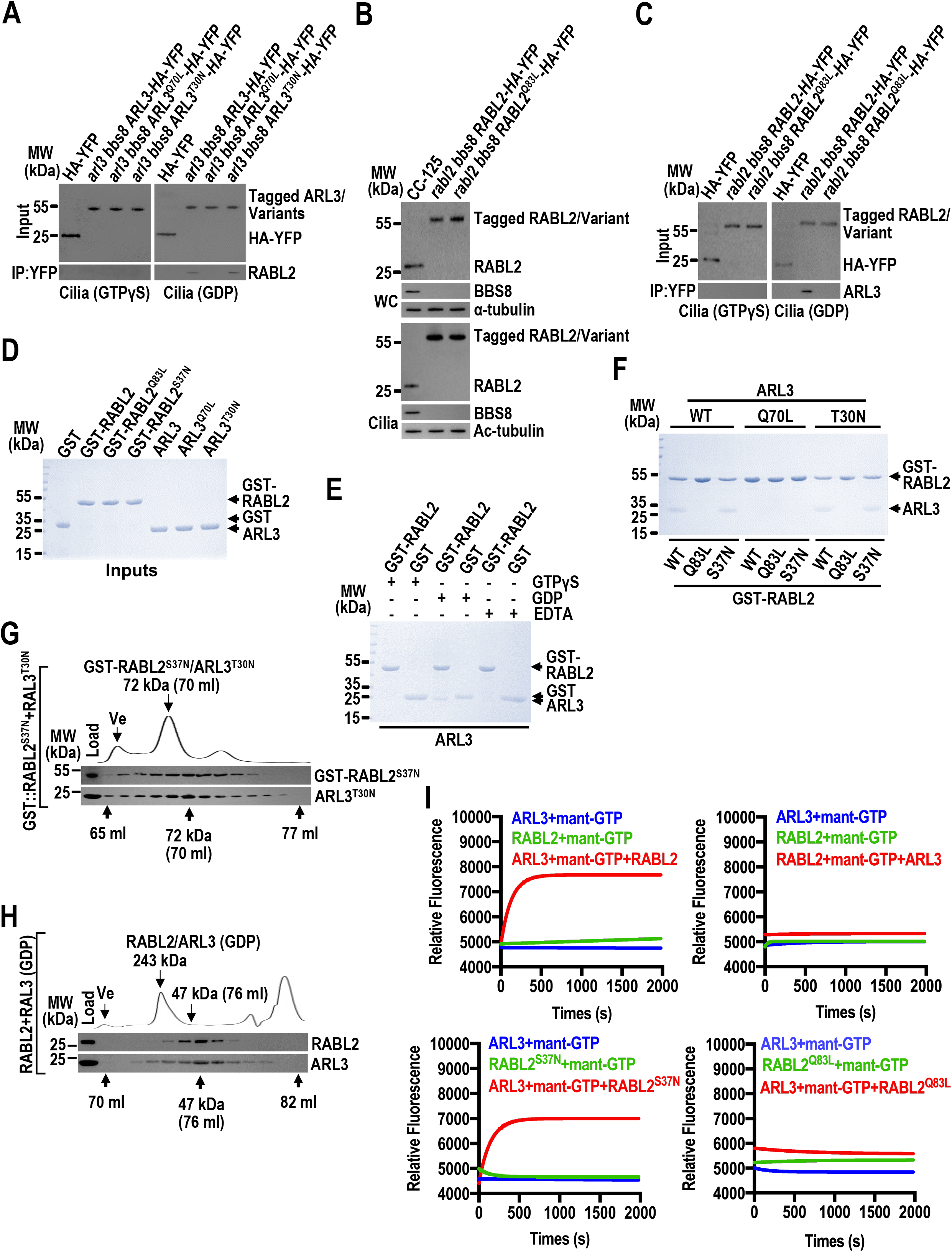
RABL2 is a ARL3-specific GEF. (*A*) Immunoblots of α-YFP-captured proteins from cilia of cells indicated on the top in the presence of GTPγS or GDP probed with α-RABL2. (*B*) Immunoblots of WC samples and cilia of cells indicated on the top probed with α-RABL2 and α-BBS8. Alpha-tubulin and acetylated (Ac)-tubulin were used as a loading control for WC samples and cilia, respectively. (*C*) Immunoblots of α-YFP-captured proteins from cilia of cells indicated on the top in the presence of GTPγS or GDP probed with α-ARL3. (*D-F*) GST, GST-RABL2, GST-RABL2^Q83L^, GST-RABL2^S37N^, ARL3, ARL3^Q70L^, and ARL3^T30N^ resolved by SDS-PAGE and visualized by Coomassie staining (*D*). GST and GST-RABL2 mixed with ARL3 in the presence of GTPγS, GDP, or EDTA and complexes recovered on glutathione beads resolved by SDS-PAGE and visualized by Coomassie staining (*E*). GST-RABL2, GST-RABL2^Q83L^, GST-RABL2^S37N^, ARL3, ARL3^Q70L^, and ARL3^T30N^ mixed to form nine combinations and complexes recovered on glutathione beads resolved by SDS-PAGE and visualized by Coomassie staining (*F*). (*G*-*H*) GST-RABL2^S37N^/ARL3^T30N^ recovered on glutathione beads (*G*) and ciliary samples of CC-125 cells (*H*) fractionated by size exclusion chromatography (S200 sizing column) and probed with α-RABL2 and α-ARL3 by immunoblotting. Relative protein concentration of the fractions between elution volume of 65 ml and 77 ml (*G*) and 70 ml and 82 ml (*H*), respectively, is shown as absorbance at 280 nm. The excluded volume (Ve) and the elution volume of GST-RABL2^S37N^/ARL3^T30N^ (72 kDa) (*G*) and RABL2/ARL3 (47 kDa) (*H*) are indicated under the immunoblots. (*I*) Mant-GTP association measurement for determining GEF activity of RABL2 on ARL3. Fluorescence intensity of mant-GTP (0.75 μm) was plotted against the recording time (s) for a combination of 2 μm ARL3, RABL2, RABL2^S37N^, or RABL2^Q83L^ alone or in the presence of 0.5 μm RABL2, RABL2^S37N^, or RABL2^Q83L^ (GEF activity on ARL3) or ARL3 (GEF activity on RABL2). Mant-GTP binding affinity values were shown. Data are averaged from three replicates. The experiment was performed three times, and one representative is shown. For panels *A* and *C*, the input was adjusted with α-YFP by immunoblotting. For panels *A*-*H*, MW stands for molecular weight.

To further discern if they bind directly, bacterially expressed N-terminal GST-tagged RABL2 (GST-RABL2) and ARL3 were generated (Fig 6*D*). In the presence of GTPγS, GDP, or EDTA that lock two GTPases simultaneously in a GTP-bound, GDP-bound, or nucleotide-free state, respectively (19), ARL3 failed to bind GST but was bound to GST-RABL2 only in the presence of GDP, confirming that two GTPases bind when they both are being GDP-locked (Fig. 6*E*). The same procedure was next performed on all combinations of wild-type, GTP-locked, or GDP-locked variants of GST-RABL2 and ARL3. They were bound to each other only when they both are under GDP-locked or wild-type conditions, revealing that bacterially expressed GTPases exists as being GDP-bound (Fig. 6*F*). They bind at a molecular ratio of 1:1 as size exclusion chromatography of the pulldown eluate showed cofractionation of GST-RABL2^S37N^ and ARL3^T30N^ with a molecular weight of 72 kDa (Fig. 6*G*). In the presence of GDP, the endogenous ARL3 and RABL2 cofractionated at a molecular weight of 47 kDa, uncovering that a transient RABL2^GDP^/ARL3^GDP^ heterodimer indeed exists in cilia (Fig. 6*H*). As determined by GEF assay, RABL2 and RABL2^S37N^ both promoted ARL3 to bind mant-GTP at a similar binding rate of 1.82×10^-4^ and 1.71×10^-4^ (μmol mant-GTP/μmol ARL3/sec), respectively (Fig. 6*I*). In contrast, RABL2^Q83L^ failed to do so (Fig. 6*I*). After the GEF effect of ARL3 on RABL2 was excluded (Fig. 6*I*), we conclude that RABL2 activates ARL3 by acting as an ARL3-specific GEF in cilia but not *vice versa*.

### RABL2^GDP^ binds ARL3^GTP^/BBSome as a BBSome cargo for outward TZ passage

Retrograde IFT train-shed RABL2^GTP^ is disabled for recoupling with IFT-B1 for ciliary retrieval via IFT as RABL2^Q83L^ fails to exit cilia but accumulates at the ciliary base right above the TZ (Fig. 3) (42). Consequently, RABL2^GTP^, once becoming autonomous of retrograde IFT train association above the TZ, must convert into RABL2^GDP^ prior to its ciliary removal (Fig. 3*G* and *SI Appendix*, Fig. S2*D*). Of note, ARL3 resided in the ciliary matrix rather than the ciliary membrane even when the BBSome was absent from cilia, revealing that RABL2^GTP^ converts into RABL2^GDP^ for activating ARL3 independent of the BBSome (Fig. 7*A*) (33). Since RABL2 relies on the BBSome for ciliary retrieval and RABL2^GTP^ fails to exit cilia even in the presence of the BBSome, we proposed that it could be RABL2^GDP^ that exits cilia via the BBSome ciliary retrieval pathway (Fig. 3 and 4).

**Figure 7.**
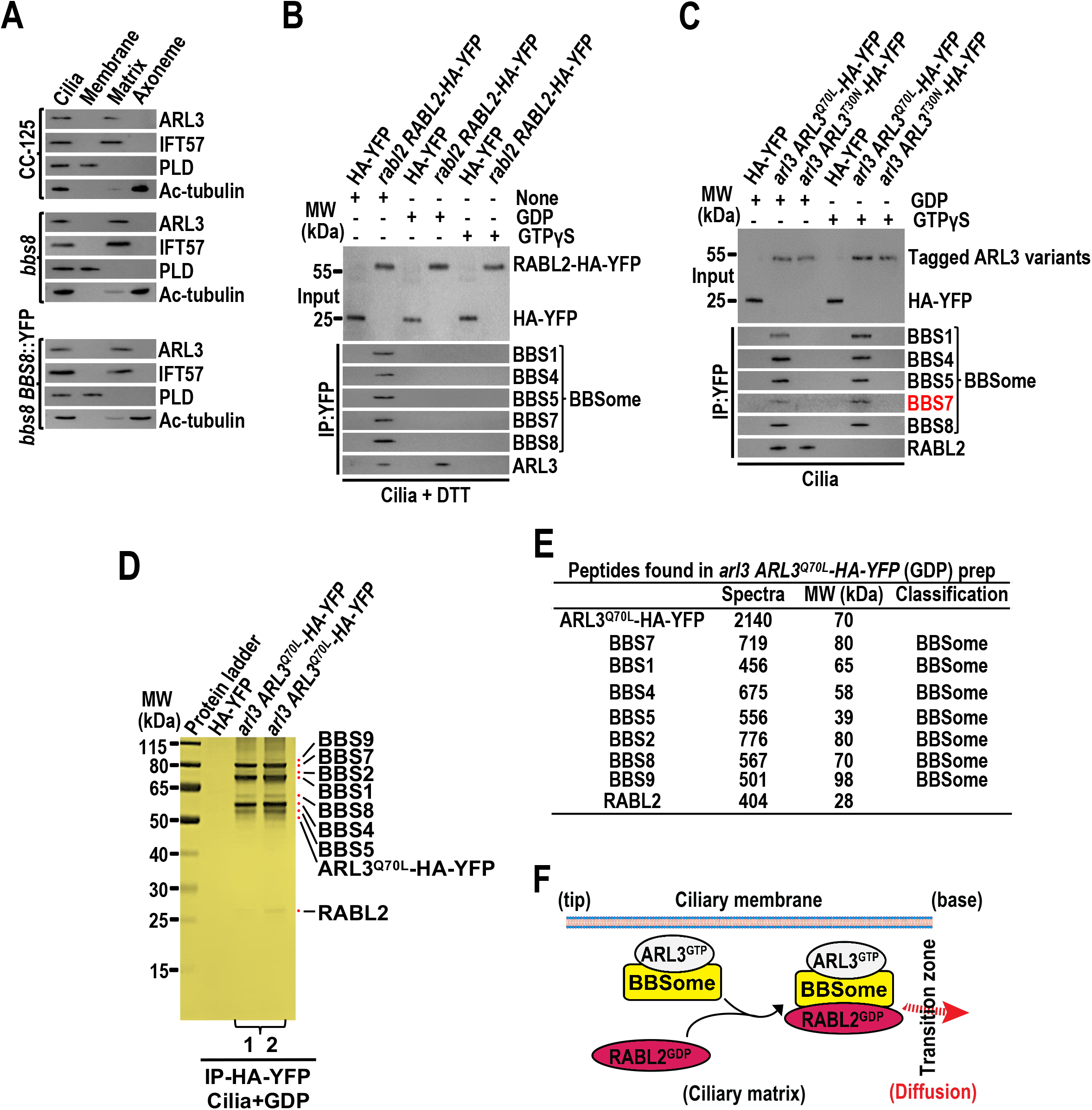
RABL2^GDP^ binds the ARL3^GTP^-bound BBSome as a cargo for passing the TZ and out of cilia. (*A*) Immunoblots of ciliary fractions of cells indicated on the left probed with α-ARL3, α-IFT57 (ciliary matrix marker), α-PLD (ciliary membrane marker), and acetylated (Ac)-tubulin (axoneme marker). (*B*) Immunoblots of α-YFP-captured proteins from cilia of cells listed on the top in the presence of GDP, GTPγS or none of them probed with α-BBS1, α-BBS4, α-BBS5, α-BBS7, α-BBS8, and α-ARL3. The assay was performed in the presence of DTT that separates the BBSome from IFT-B1. (*C*) Immunoblots of α-YFP-captured proteins from cilia of cells listed on the top in the presence of GDP or GTPγS probed with α-BBS1, α-BBS4, α-BBS5, α-BBS7, α-BBS8, and α-RABL2. (*D*) Immunoprecipitation elutes obtained from ARL3^Q70L^-HA-YFP cilia in the presence of GDP in panel *C* resolved by Bolt Bis-Tris Plus Gel and visualized by silver staining. Two folds amount of elutes was also loaded for clearly visualizing RABL2^GDP^. MW stands for molecular weight. (*E*) Tabulation of spectral counts for protein samples shown in panel *E*. (*F*) Schematic representation of how RABL2^GDP^ binds the ARL3^GTP^-bound BBSome for diffusing through the TZ for ciliary retrieval. For panels *B* and *C*, the input was adjusted with α-YFP by immunoblotting. MW stands for molecular weight.

In the presence of DTT that separates the BBSome from IFT-B1 (33), GTPγS addition failed to confer RABL2-HA-YFP to immunoprecipitate the BBSome and ARL3 (Fig. 7*B*). In the presence of GDP, RABL2-HA-YFP recovered ARL3 but not the BBSome (Fig. 7*B*). In contrast, they both were recovered by RABL2-HA-YFP when none of GDP and GTPγS was added (Fig. 7*B*). RABL2^GTP^ and RABL2^GDP^ then did not interact with ARL3^GTP^-bound and ARL3-free BBSome, respectively. It is RABL2^GDP^ that bind ARL3^GTP^-bound BBSomes. Supportively, ARL3^Q70L^-HA-YFP immunoprecipitated both the BBSome and RABL2 only when RABL2 was locked in a GDP-bound state by GDP addition (Fig. 7*C*). ARL3^T30N^-HA-YFP failed to recover the BBSome and RABL2 in the presence of GTPγS but recovered RABL2 alone when GDP was present (Fig. 7*C*). As identified by “in-solution” mass spectrometry analysis of immunoprecipitation elutes from cilia in the presence of GDP, ARL3^Q70L^-HA-YFP recovered all seven BBSome subunits and RABL2^GDP^ as well (Fig. 7*D* and *E*). We are unable to visualize RABL2 for deciding if it diffuses through the TZ for ciliary retrieval (Fig. 3*B* and *SI Appendix*, Fig. S2*B* and *C*). However, RABL2^GDP^ binds ARL3^GTP^/BBSomes as a BBSome cargo, suggesting that RABL2^GDP^ diffuses through the TZ for ciliary retrieval via the ARL3^GTP^/BBSome pathway (Fig. 7*F*) (33).

### RABL2 mediates phototaxis via the ARL3^GTP^/BBSome pathway

The loss of ARL3 impairs BBSome ciliary turnover for causing the generation of *Chlamydomonas* cells defective in phototaxis (18, 39). Since RABL2 maintains BBSome ciliary turnover via ARL3, its depletion supposedly causes the similar non-phototactic phenotype (33). Unfortunately, *rabl2^CLiP^* and *rabl2 RABL2^S37N^-HA-YFP* cells are bald (Fig. 8*A* and *SI Appendix*, Fig. S1*C*). *rabl2 RABL2-HA-YFP, rabl2 RABL2^D76G^-HA-YFP*, and *rabl2 RABL2^Q83L^-HA-YFP* cells are normal in ciliation, while they were defective in phototaxis as they are derived from the phototaxis-deficient CC-5325 cells (Fig. 8*A* and *SI Appendix*, Fig. S1*C* and S3*C*) (33). We then chosen the CC-125-derived RABL2^Res-WT^, RABL2^Res-D76G^, and RABL2^Res-Q83L^ cells for testing the role of RABL2 in mediating phototaxis (Fig. 1*F* and *G*). These cells retained cellular BBSome proteins at WT levels (Fig. 8*B*). Consistent with the RABL2 knockout and rescue cells, RABL2^Res-D76G^ and RABL2^Res-Q83L^ differed from RABL2^Res-WT^ in that they accumulated the BBSome (represented by BBS8) at the ciliary base above the basal bodies (double labeled by IFT81 and BBS8) (Fig. 8*B* and *C*). As determined by both population and single cell locomotion assays, RABL2^miRNA^ cells are defective in phototaxis, RABL2^Res-WT^ cells, like WT CC-125 cells, become normal in phototaxis, and RABL2^Res-D76G^ and RABL2^Res-Q83L^ cells remain to be defective in phototaxis (Fig. 8*D*). Therefore, RABL2 regulates phototaxis of *Chlamydomonas* cells via the ARL3^GTP^/BBSome pathway (Fig. 8*E*).

**Figure 8.**
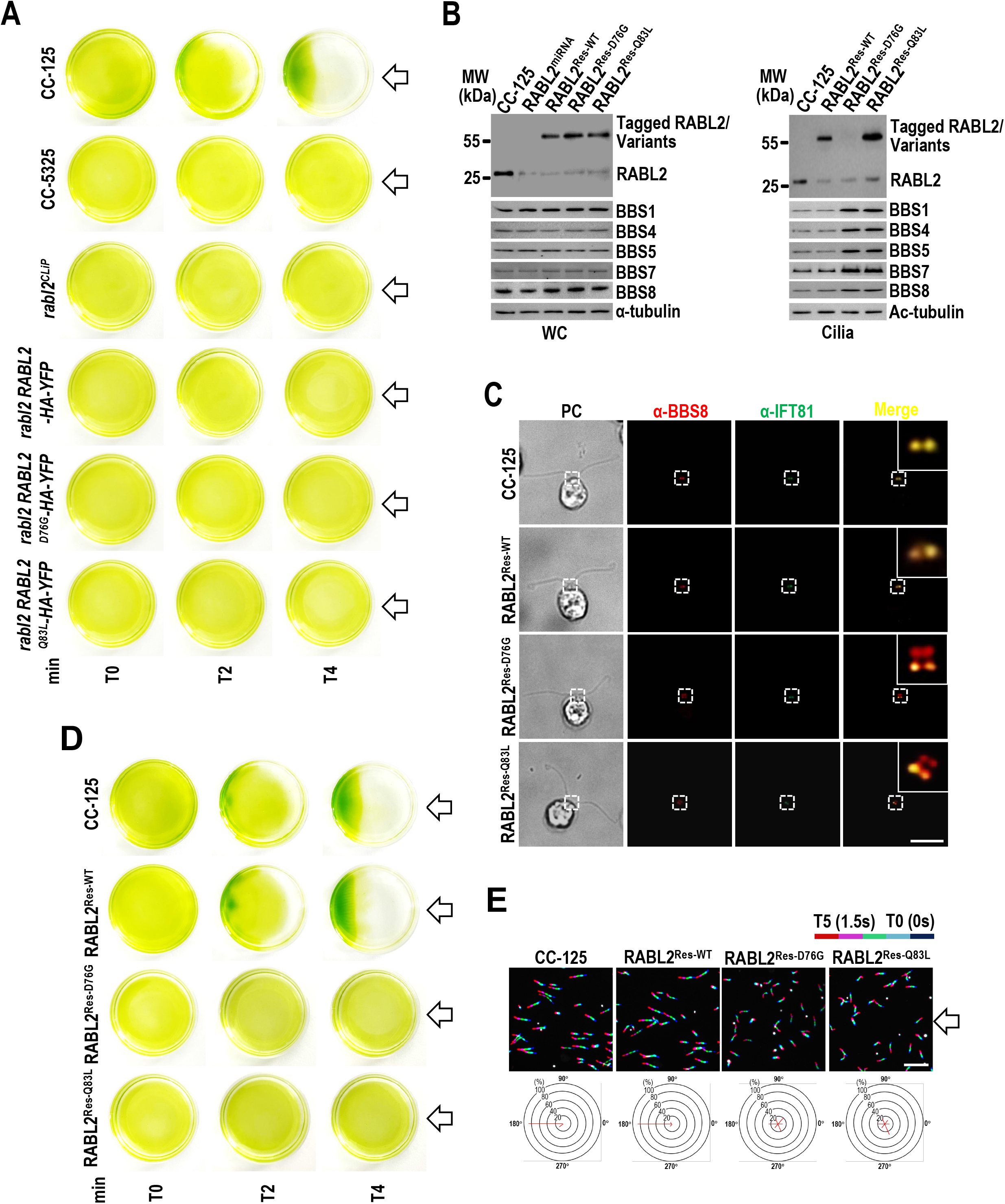
RABL2 mediates phototaxis via the ARL3/BBSome pathway. (*A*) Population phototaxis assay of cells listed on the left. (*B*) Immunoblots of WC samples and cilia of cells listed on the top probed with α-RABL2, α-BBS1, α-BBS4, α-BBS5, α-BBS7, and α-BBS8. Alpha-tubulin and acetylated (Ac)-tubulin were used for adjusting the loading for WC samples and cilia, respectively. MW stands for molecular weight. (*C*) Cells listed on the left stained with α-BBS8 (red) and α-IFT81 (green). Inset shows the basal body and the transition zone staining. PC images of cells were shown. Scale bars: 10 μm. (*D*) Population phototaxis assay of cells listed on the left. (*E*) Single cell locomotion analysis of cells listed on the top. The radial histograms show the percentage of cells moving in a particular direction relative to the light (six bins of 60°each). Composite micrographs showing the tracks of single cells. Each of the five merged frames was assigned a different color (blue, frame 1, and red, frame 5, corresponding to a travel time of 1.5 s). (Scale bar: 50 μm.). For panels *A*, *D*, and *E*, the direction of light is indicated (white arrows, flashlight).

## Discussion

We used *C. reinhardtii* as a model organism and observed that RABL2 recruits IFT-B1 as its effector to the basal bodies, deciding its basal body amount available for assembling anterograde IFT trains (Fig. 9). RABL2^GTP^ cycles through cilia as an IFT-B1 cargo (Fig. 9). Upon arriving at the ciliary base right above the TZ following retrograde transportation, RABL2^GTP^ sheds from retrograde IFT trains prior to a rapid GTP hydrolysis. RABL2^GDP^ then activates the ciliary membrane anchored ARL3^GDP^ to become ARL3^GTP^ as an ARL3-specific GEF. The active ARL3^GTP^ dissociates with the ciliary membrane and recruits the retrograde IFT train-shed and cargo (e.g. PLD)-laden BBSome as its effector to diffuse through the TZ for ciliary retrieval (Fig. 9) (33). RABL2^GDP^ itself pass the TZ and out of cilia via binding the ARL3^GTP^-bound BBSome as its cargo. RABL2 mediates anterograde IFT train assembly at the basal bodies and outward TZ passage of the BBSome via ARL3, closing a gap in our understanding of how RABL2 plays multifaceted roles both inside and outside cilia to affect cell behaviors, e.g., ciliation and phototaxis, of *C. reinhardtii*.

**Figure 9.**
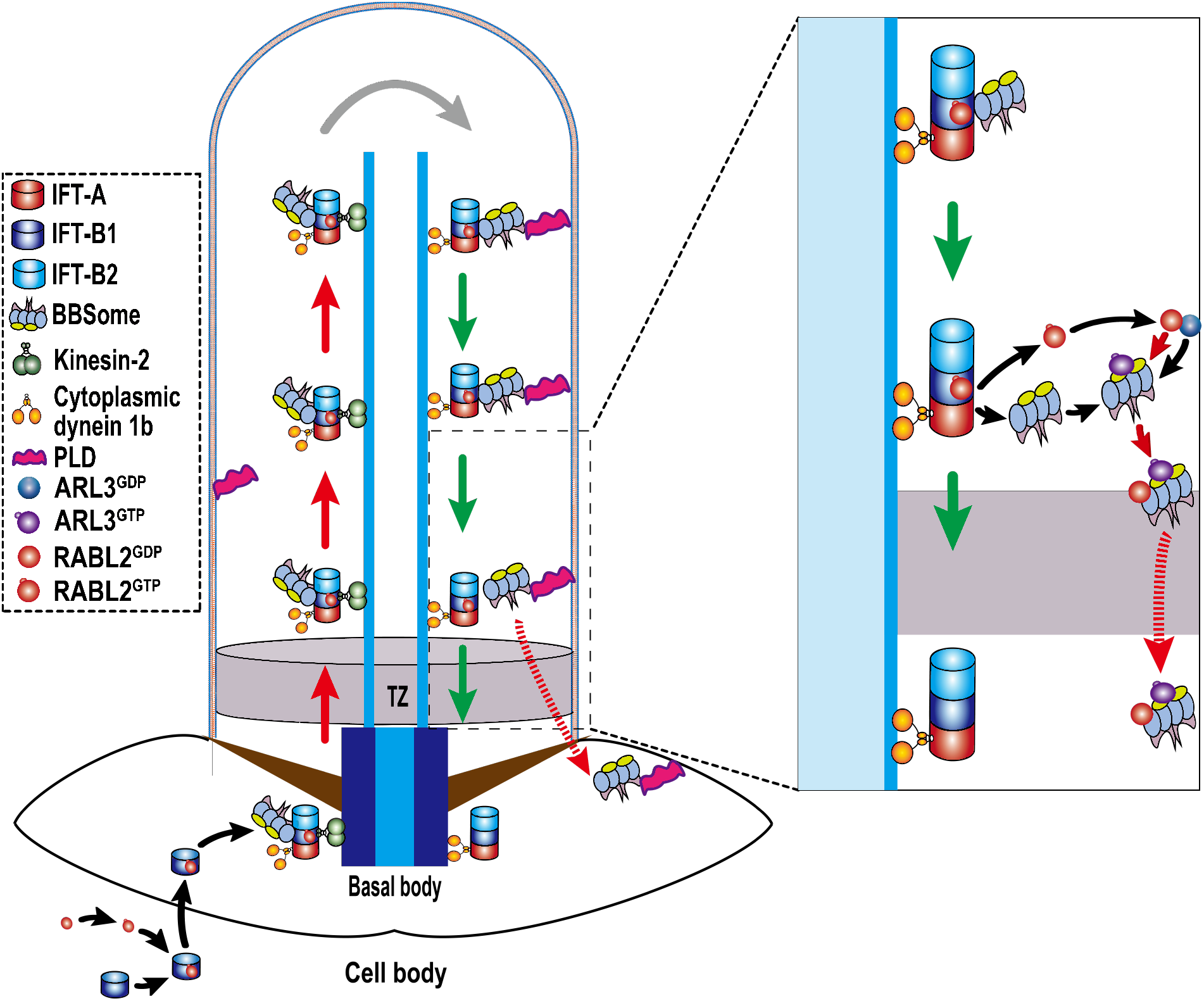
Hypothetical model of how RABL2 mediates IFT-B1 basal body recruitment and BBSome ciliary retrieval via ARL3 in *C. reinhardtii*. RABL2 exists as a GDP-bound state (RABL2^GDP^) in cytoplasm and can convert to RABL2^GTP^ intrinsically (43). RABL2^GTP^ binds and recruits its IFT-B1 effector to the basal bodies. At the basal bodies, IFT-B1 integrates into anterograde IFT trains for initiating ciliation. RABL2^GTP^ binds IFT-B1 for cycling through cilia via IFT. Upon reaching the proximal ciliary region right above the TZ in the retrograde direction, RABL2^GTP^ sheds from IFT-B1 followed by hydrolyzing to become RABL2^GDP^. At the same ciliary region, ciliary membrane-anchored ARL3^GDP^ is bound to and activated by RABL2^GDP^ as a ARL3-specific GEF. Following the transportation from the ciliary tip to base, the signaling molecule (e.g. PLD)-laden BBSome sheds from retrograde IFT trains right above the TZ region and is bound to ARL3^GTP^ as an ARL3-specific effector (33). ARL3^GTP^ then recruits the PLD-laden BBSome to diffuse through the TZ for ciliary retrieval (33). RABL2^GDP^ binds the ARL3^GTP^-bound BBSome as a BBSome cargo for diffusing through the TZ for ciliary retrieval.

### How does RABL2 mediate ciliation?

RABL2 uses IFT-B1 as its effector in human and murine cells (42–45). RABL2 resides at the ciliary base in the steady state, while human RABL2 rather than the murine orthologue controls ciliation as shown in RPE-hTERT cells (42–44). *Chlamydomonas* RABL2 resembles the human and murine orthologues in using IFT-B1 as its effector and recruits IFT-B1 to the basal bodies, mediating ciliation by regulating IFT-B1 basal body amount available for assembly of anterograde IFT trains (Fig. 2 and 9). Interestingly, knockout of murine RABL2 does not disrupt IFT and ciliation, suggesting that murine RABL2 simply binds IFT-B1 as a cargo for trafficking to the basal bodies so that it can enter and cycle through cilia via IFT (42). In this case, certain GTPase(s) other than RABL2 may regulate IFT-B1 targeting to the basal body as ciliation has been shown to rely on the integrity of anterograde IFT trains assembled at the basal bodies (43, 50). Our previous study and others have identified RABL5/IFT22 and ARL6/BBS3 coordinate to recruit the BBSome to the basal bodies, controlling its amount available for integrating into anterograde IFT trains as an IFT cargo adaptor (21, 40). Together with the current finding, we propose that cells may apply GTPases as molecular switches for recruiting the IFT-A and -B2 component subcomplexes of anterograde IFT trains to the basal bodies. By such a strategy, cells are supposed to regulate the basal body amounts of all three IFT subcomplex components available for assembling anterograde IFT trains for initiating IFT. It is also worth noting that the RABL2^D76G^ mutant enables IFT-B1 to target to the basal bodies but does not enter cilia itself, revealing that IFT and ciliation likely remain to be normal even when RABL2 is not present in cilia (Fig. 4). As compared to IFT proteins that are easily visualized to reside at the basal bodies and in cilia, RABL2 even tagged with super bright fluorescent proteins such as 3NG is undetectable through fluorescent microscopy, revealing that only a very small portion of IFT-B1 entering cilia is RABL2-loaded (Fig. 3*B* and *E* and *SI Appendix*, Fig. S2*C*). For this reason, human RABL2 still could enter hTERT-RPE cilia (43). RABL2 ciliary absence in human cells may reflect its extremely low ciliary abundance, which hardly can be easily detected by regular immunostaining (43).

### How does RABL2 activate ARL3 for recruiting the BBSome to diffuse through the TZ for ciliary retrieval?

RABL2 and ARL3 bind one another in a molecular ratio of 1:1 only when they both are in a GDP-bound state (Fig. 6*D-G*). Indeed, RABL2^GDP^ activates ARL3^GDP^ to become ARL3^GTP^ but not *vice versa*, revealing that RABL2 acts as an ARL3-specific GEF (Fig. 6*I*). As expected, transient interaction of RABL2 with ARL3 was captured to exist in cilia only in the presence of GDP that locks both GTPases in a GDP-bound state (Fig. 6*H*). Our previous study has identified ARL3^GDP^ associates with the membrane for diffusing into cilia under physiological conditions (19). Upon inside cilia, ARL3^GDP^ is rapidly converted to ARL3^GTP^ by an unknown mechanism (19). Following this activation process, ARL3^GTP^ dissociates with the ciliary membrane and recruits the retrograde IFT train-shed and cargo (e.g. PLD) laden BBSome as its effector to diffuse through the TZ for ciliary retrieval (19). Our current study provided compelling functional and biochemical evidence to show that it is RABL2 that activates ARL3 for recruiting the BBSome to diffuse through the TZ for ciliary retrieval. According to our current data, we propose that RABL2^GTP^ enters and cycles through cilia by acting as an IFT-B1 cargo (Fig. 2 and 3). Upon at the proximal ciliary region right above the TZ following retrograde IFT, RABL2^GTP^ sheds from IFT-B1 followed by converting into RABL2^GDP^, which binds and activates the ciliary membrane anchored ARL3^GDP^ as an ARL3-specific GEF (Fig. 9). Afterward, ARL3^GTP^ dissociates with the ciliary membrane and recruits the retrograde IFT train-shed and cargo (e.g. PLD)-laden BBSome to diffuse through the TZ for ciliary retrieval (Fig. 9) (33).

### Howe does RABL2 cycle through cilia?

Except for human RABL2 that fails to enter cilia, murine and *Chlamydomonas* RABL2^GTP^ binds IFT-B1 as a cargo protein for entering and cycling cilia via IFT (Fig. 3) (42, 43). Murine RABL2 was proposed to hydrolyze GTP for converting into RABL2^GDP^ so as to be able to jump off the IFT-B1 complex component of retrograde IFT at the proximal ciliary region right above the TZ (42). Our data cannot exclude this possibility from happening in *C. reinhardtii*, while GTP-locked RABL2^Q83L^ deprived of GTP hydrolysis continues to shed from retrograde IFT trains right above the TZ region, supporting the notion that RABL2^GTP^ can jump off IFT-B1 prior to its conversion into RABL2^GDP^ (Fig. 3*E* and *G*) (43). It is also worth noting that GTP-bound RABL4/IFT27 by binding IFT25 to form a IFT25/27 heterodimer can jump on and off the IFT-B1 subcomplex component of IFT trains at the ciliary tip, suggesting that certain Rab-like GTPases do not rely on their nucleotide state for coupling/uncoupling with their “effectors” in these specialized ciliary compartments (e.g. the ciliary tip and base) (19). However, RABL2 differs from IFT27 in that IFT-B1-shed RABL2^GTP^ is disabled for reloading onto IFT-B1 for ciliary retrieval via IFT. That is why RABL2^Q83L^ fails to exit cilia but accumulates at the proximal ciliary region right above the TZ (Fig. 3*E*). At present, the mechanism behind how RABL2^GTP^ converts into RABL2^GDP^ at the ciliary base remains unknown (Fig. 3*G* and *SI Appendix*, Fig. S2*D*). However, *Chlamydomonas* RABL2 was shown to has intrinsic GTPase activity, a clue excluding its nucleotide exchange from requiring a GEF (43). Upon activating ARL3, RABL2^GDP^ specifically binds ARL3^GTP^/BBSome, autonomous of retrograde IFT train association, as a BBSome cargo followed by diffusing through the TZ for ciliary retrieval via the ARL3^GTP^/BBSome ciliary retrieval pathway, consistent with the observation that RABL2 and the BBSome relies on one another for passing the TZ and out of cilia (Fig. 4). Therefore, RABL2 enters cilia being an IFT-B1 cargo and exits cilia as a BBSome cargo. During these processes, a RABL2 GTPase cycle is finished in cilia.

### Implications for the molecular base of bardet-biedl and Joubert syndromes

Murine RABL2^D73GQ80L^ double mutant (RABL2^D76GQ83L^ in *Chlamydomonas* accordingly) resembles RABL2 to uncouple with retrograde IFT at the ciliary base so as to promote outward TZ passage of the BBSome (42). When expressed at the endogenous RABL2 level in *rabl2^CLiP^* (resulting strain *rabl2* RABL2^D76GQ83L^-HA-YFP), RABL2^D76GQ83L^-HA-YFP accumulated in cilia to an extent even more than RABL2^Q83L^-HA-YFP (*SI Appendix*, Fig. S5*A*). RABL2^D76GQ83L^-HA-YFP resembled RABL2^Q83L^-HA-YFP in performing normal IFT but accumulated at the CEP290-labled TZ region, indicating that its ciliary retention is caused by its inability to pass the TZ for ciliary retrieval (*SI Appendix*, Fig. S5*B-E*). Therefore, D76G and Q83L double mutations do not disrupt RABL2 uncoupling with retrograde IFT at the ciliary base. RABL2^D76GQ83L^ is supposed to be disabled for converting to being GDP-bound. This hampered ARL3 activation and prevented the BBSome (represented by BBS8) from passing the TZ for ciliary retrieval but accumulated at the ciliary base right above the TZ by co-localizing with RABL2^D76GQ83L^-HA-YFP (*SI Appendix*, Fig. S5*F*). These RABL2 variants were not identified for causing BBS in humans thus far, while mice carrying these mutations resemble the IFT27 knockout for displaying similar symptoms (42, 45, 49). As IFT27 is a regulator of BBSome function and implicated for causing BBS (*BBS19*) (51), RABL2 dysfunction is expected to cause BBS. In addition, ARL3 mutations cause JS in humans (52). According to the functional overlap of two small GTPase over BBSome signaling, RABL2 dysfunction is also expected to cause JS.

## Materials and methods

### Antibodies, *Chlamydomonas* strains, and culture conditions

Rabbit-raised polyclonal antibodies against IFT38, IFT43, IFT46, IFT57, IFT70, IFT139, BBS1, BBS4, BBS5, BBS7, BBS8, CEP290, ARL3, and PLD have been reported previously (39–41, 53). Antibodies against YFP (mAbs 7.1 and 13.1, Roche), α-tubulin (mAb B512, Sigma-Aldrich) and acetylated-α-tubulin (mAb 6-11B-1, Sigma-Aldrich) were commercially bought. Rabbit-raised polyclonal antibody against RABL2 was made by Beijing Protein Innovation LLC (Beijing, China). For immunoblotting and immunostaining assays, antibodies were diluted properly as listed in *SI Appendix*, Table S1. *Chlamydomonas* strain CC-125 was purchased from the *Chlamydomonas* Resource Center at the University of Minnesota, Twin Cities, MN (http://www.chlamycollection.org/). CC-5325, *arl3^CLiP^* (LMJ.RY0420.182282), and *rabl2^CLiP^* (LMJ.RY0402.205222) were available from the *Chlamydomonas* Library Project (CLiP, https://www.chlamylibrary.org/allMutants) (54). Strains *bbs8* (55), *ift46-1* IFT46-YFP (48), HA-YFP (47), *arl3 ARL3-HA-YFP, arl3 ARL3^Q70L^-HA-YFP, arl3 ARL3^T30N^-HA-YFP, arl3 bbs8 ARL3-HA-YFP*, *arl3 bbs8 ARL3^Q70L^-HA-YFP*, and *arl3 bbs8 ARL3^T30N^-HA-YFP* (33) have been reported previously. All the strains used in this study were listed in *SI Appendix*, Table S2. *Chlamydomonas* strains were cultured at room temperature in Tris acetic acid phosphate (TAP) medium or minimal 1 (M1) medium in a continuous light with constant aeration. Depending on a specific strain, cells were cultured with or without the addition of 20 μg/ml paromomycin (Sigma-Aldrich), 15 μg/ml bleomycin (Invitrogen) or both antibiotics with 10 μg/ml paromomycin and 5 μg/ml bleomycin.

Various experimental protocols were applied in this study, and most of the experiments were briefly introduced in the text to make the content easier to be understood. The details of each protocol are available in *SI Appendix*.

### Data availability

Data supporting the findings of this study were contained within this paper and SI Appendix.

## Acknowledgements

Research reported in this publication was supported by National Natural Science Foundation of China (32070698 to Z-C.F.) and China Postdoctoral Science Foundation (2021M702457 to Y-X.L.). The founder has no role in study design, data collection and analysis, decision to publish, or preparation of the manuscript.

## Author contributions

Z-C.F. directed and designed the study. R-K.Z., W-Y.S. and Y-X.L. performed functional and biochemical validation experiments. R-K.Z., W-Y.S. and Y-X.L. performed immunostaining assays. Y-X.L. and R-K.Z. performed GEF and silver staining assays and mass spectrometry data analysis. R-K.Z. performed TIRF imaging of living cells. R-K.Z. performed transgenic *Chlamydomonas* strain screening. D-X.B., R-Q.J. and C.Z. helped to screen transgenic *Chlamydomonas* cells. Z-C.F. analyzed the data and wrote the manuscript.

