## Supplemental materials for "RABL2 Regulates Ciliation via Controlling IFT-B1 Basal Body Recruitment and ARL3-mediated BBSome Ciliary Retrieval"

**This PDF file includes:**

Materials and methods
Figures S1 to S5
Tables S1 to S3
Legends for Movies S1 to S10
SI References

**Other supplementary materials for this manuscript include the following:**

Movies S1 to S10

### Materials and methods

**Plasmids and strain generation.** RABL2 miRNA vector was generated according to the method described previously (1). In brief, the miRNA sequence targeting the 3'-UTR region of RABL2 gene was designed using WMD3 software (http://wmd3.weigelworld.org) and was combined with the miRNA cre-MIR1157 (accession number MI0006219) to result in a 171 bp of RABL2 miRNA precursor sequence (RABL2-miRNA-3'-UTR) (SI Appendix, Table S3). The miRNA precursor sequence was synthesized by Genewiz (China) and ligated to the pHK263 plasmid (1), resulting in the RABL2 miRNA vector pMi-RABL2-Paro. *Chlamydomonas* expression vectors were generated on a pBluescript II-KS+ backbone and contained hemagglutinin (HA) and yellow fluorescent protein (YFP) (HA-YFP) coding sequences followed immediately downstream by a sequence encoding the Rubisco 3'-UTR and the *aph*VIII cassette (paromomycin resistant gene), the *ble* cassette (zeocin resistant gene), or the *hyg* cassette (hygromycin B resistant gene).

To express RABL2-HA-YFP, a 3,241-bp genomic DNA fragment composed of 1,091-bp promoter sequence and the coding region of RABL2 was amplified from genomic DNA by using the primer pair gRABL2-FOR and gRABL2-REV as listed in SI Appendix, Table S3, double digested with *Xba*I and *Eco*RI, and inserted into pBluescript II KS(+), resulting in pBKS-gRABL2. Afterwards, the gRABL2 sequence cut from pBKS-gRABL2 by double digestion with *Xba*I and *Eco*RI and the HA-YFP-ble sequence cut from pBKS-gBBS3-HA-YFP-Ble by double digestion with *Eco*RI and *Kpn*I (2) were inserted into the *Xba*I and *Kpn*I sites of pBluescript II KS(+) vector, resulting in pBKS-gRABL2-HA-YFP-Ble.

To express RABL2$^{Q83L}$-HA-YFP, RABL2$^{S37N}$-HA-YFP, RABL2$^{D76G}$-HA-YFP, and RABL2$^{D76GQ83L}$-HA-YFP, the desiring mutations were introduced into pBKS-gRABL2 by site-directed mutagenesis using the primer pairs (RABL2$^{Q83L}$-FOR and RABL2$^{Q83L}$-REV, RABL2$^{S37N}$-FOR and RABL2$^{S37N}$-REV, RABL2$^{D76G}$-FOR and RABL2$^{D76G}$-REV) as listed in SI Appendix, Table S3, resulting in pBKS-gRABL2$^{Q83L}$, pBKS-gRABL2$^{S37N}$, pBKS-gRABL2$^{D76G}$ and pBKS-gRABL2$^{D76GQ83L}$. Same cloning strategy as shown above was used to generate pBKS-gRABL2$^{Q83L}$-HA-YFP-Ble, pBKS-gRABL2$^{S37N}$-HA-YFP-Ble, pBKS-gRABL2$^{D76G}$-HA-YFP-Ble and pBKS-gRABL2$^{D76GQ83L}$-HA-YFP-Ble.

To express RABL2-HA-NG, the HA-NG sequence was amplified from pLV-mNeonGreen (3) by using primer pair (HA-NG-FOR and HA-NG-REV) as listed in SI Appendix, Table S3, double digested with *Eco*RI and *Xho*I, and inserted into pBKS-gRABL2-HA-YFP-Ble to replace the HA-YFP fragment, resulting in pBKS-gRABL2-HA-NG-Ble. To express RABL2-HA-3NG, Three NG fragments that we named NG-1, NG-2, and NG-3 were amplified from pBKS-gRABL2-HA-NG-Ble by using primer pairs (NG-1-FOR and NG-1-REV, NG-2-FOR and NG-2-REV, NG-3-FOR and NG-3-REV) as listed in SI Appendix, Table S3. NG-1 and NG-2 fragments were double digested with *Eco*RI/*Bam*HI and *Bam*HI/*Hind*Ⅲ, respectively and inserted into the *Eco*RI and *Hind*III sites of pBluescript II KS(+) by three-way ligation, resulting in pBKS-HA-NG-1-NG-2. Next, the NG-3 fragment was double digested with *Hin*dIII and *Xho*I and inserted into pBKS-HA-NG-1-NG-2, resulting in pBKS-HA-3NG. The HA-NG fragment of pBKS-gRABL2-HA-NG-Ble was replaced with the HA-3NG fragment using the same cloning strategy as shown above, resulting in pBKS-gRABL2-HA-3NG-Ble.

To express RABL2-HA, RABL2$^{Q83L}$-HA, RABL2$^{D76G}$-HA and RABL2$^{D76GQ83L}$-HA, the gRABL2-HA, gRABL2$^{Q83L}$-HA, gRABL2$^{D76G}$-HA, and gRABL2$^{D76GQ83L}$-HA fragments were amplified from pBKS-gRABL2, pBKS-gRABL2$^{Q83L}$, pBKS-gRABL2$^{D76G}$, and pBKS-gRABL2$^{D76GQ83L}$ by using primer pair (gRABL2-FOR and gRABL2-HA-REV) as listed in SI Appendix, Table S3. Afterwards, these amplified DNA fragments digested with *Xba*I and *Xho*I sites and the Ble sequence derived from pBKS-gRABL2-HA-YFP-Ble by digestion with *Xho*I and *Kpn*I were inserted into *Xba*I and *Kpn*I sites of pBluescript II KS(+) vector by three-way ligation, resulting in pBKS-gRABL2-HA-Ble, pBKS-gRABL2$^{Q83L}$-HA-Ble, pBKS-gRABL2$^{D76G}$-HA-Ble, and pBKS-gRABL2$^{D76GQ83L}$-HA-Ble.

To express ARL3$^{Q70L}$-HA-YFP in *rabl2*::RABL2$^{Q83L}$-HA-YFP strain , HA-YFP sequence was cut from pBKS-gRABL2-HA-YFP-Ble by *Eco*RI and *Apa*I, and inserted into the *Eco*RI and *Apa*I sites of pBluescript II KS(+) vector, resulting in pBKS-HA-YFP. Next, HYG sequence was inserted into *Kpn*I site of pBKS-HA::YFP by the recombinase, resulting in pBKS-HA-YFP-Hyg. Afterwards, a 3,342-bp ARL3$^{Q70L}$ fragment consisting of the 1,066-bp promoter sequence and its coding region was amplified from pBKS-gARL3$^{Q70L}$-HA-YFP-Ble by using primer pair (gARL3$^{Q70L}$-FOR and gARL3$^{Q70L}$-REV) as listed in SI Appendix, Table S3 (4), gARL3$^{Q70L}$ was inserted into *Xba*I site of pBKS-HA::YFP-Hyg by the recombinase, resulting in pBKS-gARL3$^{Q70L}$-HA-YFP-Hyg.

The new constructs were verified by direct nucleotide sequencing before they were transformed into *Chlamydomonas* cells by electroporation as described previously and screening of the positive transformants was done according to the method described previously (5). The screening of RABL2 miRNA cells was initiated by checking the cellular level of the target proteins through immunoblotting of whole cell extracts with RABL2 antibody. The miRNA strain showing a reduced level of RABL2 was selected for further phenotypic analysis. Double mutant generation has been described previously (6).

**Preparation of cell bodies and cilia.** Cilia and cell bodies were prepared as described in our previous report (7). Briefly, 5 liters of *Chlamydomonas* cells were centrifuged at 1,000 ×*g* at 4°C for 15 min for collection followed by suspending in 150 ml of TAP (pH7.4) and incubating for 2 hrs under strong light with continuous bubbling. Afterwards, 0.5 M acetic acid was added to reach a pH value of 4.5 for deciliating cells before 0.5 M KOH was added to reach a pH value of 7.4. The solution was centrifuged at 600 ×*g* at 4°C for 5 min for separating cell bodies (pellets) from cilia (supernatants). Cilia were then washed with HMDEKN buffer (30 mM Hepes, [pH 7.4], 5 mM

MgSO4, 1 mM dithiothreitol [DTT], 5 mM EGTA, and 25 mM KCl, 125 mM NaCl) by centrifugation at 12,000 ×*g* at 4°C for 10 min until no green color observed.

**Preparation of ciliary fractions.** Ciliary fractions were prepared as described in our previous report (2). In brief, cilia were first dissolved in HMDEKN buffer and protein inhibitors (PI) (1 mM PMSF, 50 μg/ml soybean trypsin inhibitor, 1 μg/ml pepstatin A, 2 μg/ml aprotinin, and 1 μg/ml leupeptin) and fresh-frozen in liquid nitrogen. After three frozen-and-thaw cycles, the ciliary matrix fraction was obtained by centrifugation at 12,000 ×g at 4 °C for 15 min. The pellets were dissolved in HMDEKN buffer containing 0.5% NP-40 and stayed on ice for 15 min before centrifuged at 12,000 ×*g* at 4°C for 10 min. After centrifugation, the supernatant and pellet were collected as the membrane and axonemal fractions, respectively.

**Immunoblotting.** Whole cell, cell body, and ciliary extracts were prepared as described in our previous report (8). SDS-PAGE electrophoresis, PVDF membrane transfer, and immunoblotting were performed as previously described (9). If not otherwise specified, 20 µg of proteins from each sample was loaded for SDS-PAGE electrophoresis. The primary and secondary antibodies were diluted for immunoblotting with a ratio as shown in SI Appendix, Table S1. If applicable, the target proteins were quantified by measuring the intensity of the immunoblots with ImageJ software (version 1.42g, National Institutes of Health) as described previously (5). The intensity of immunoblots was normalized to the intensity of a loading control protein and presented using GraphPad Prism 8.0 (GraphPad Software).

**Immunoprecipitation.** Immunoprecipitation was performed as described in our previous report (10). Briefly, cell bodies and cilia isolated from *Chlamydomonas* cells expressing HA-YFP- or HA-NG-tagged bait proteins were resuspended in HMEKN buffer (30 mM Hepes, [pH 7.4], 5 mM MgSO4, 5 mM EGTA, and 25 mM KCl, 125 mM NaCl) plus PI followed by adding NP-40 to 1% to lyse the cell. The supernatants were collected by centrifugation at 14,000 ×*g* for 10 min and were incubated with agitation with 5% BSA-pretreated camel anti-GFP or anti-NG antibody-conjugated agarose beads (V-nanoab Biotechnology) for 2 hrs. By centrifugation at 2,500 ×*g* for 5 min, the beads were washed for three times with HMEKN buffer. The beads were then added with Laemmli SDS sample buffer and boiled for 5 min before centrifugation at 2,500 ×*g* for 5 min. The supernatants were then collected for analyzing by immunoblotting. The centrifugation was done at 4°C. If necessary, immunoprecipitation was performed in the presence of GTPγS (20 mM) or GDP (20 mM).

**Immunofluorescence.** Immunofluorescence staining of *Chlamydomonas* cells was performed as described in our previous report (9). The primary antibodies including rabbit-raised α-CEP290, α-

BBS8, α-IFT38, α-IFT43, α-IFT46, and α-IFT70, and mouse-raised α-IFT81 and α-YFP, and the secondary antibodies Alexa-Fluor594 conjugated goat anti-rabbit IgG, and Alexa-Fluor488 conjugated goat anti-mouse IgG, and Alexa-Fluor594 conjugated goat anti-mouse IgG (Molecular Probes) were listed in SI Appendix, Table S1 with their suggested dilutions for immunostaining. Images were acquired with an IX83 inverted fluorescent microscope (Olympus) equipped with a back illuminated scientific CMOS camera (Prime 95B, Photometrics), a 100×/1.40 NA oil objective lens (Olympus), and 488-nm and 561-nm lasers from Coherent OBIS Laser Module. All images were captured and processed with CellSens Dimension (version 2.1, Olympus).

**Sucrose density gradient centrifugation.** Sucrose density gradient centrifugation was performed on cell body and ciliary extracts according to our published method (8). Briefly, linear 12 ml of 10-25% sucrose density gradients in 1×HMDEKN buffer plus PI and 1% NP-40 were generated. Cell body and cilia were treated for frozen-and-thaw by liquid nitrogen and centrifuged at 12,000 ×*g* at 4 °C for 10 min. Seven hundred micro-litter of cell body or ciliary extracts were loaded on the top of the gradients and separated at 38,000 ×*g* at 4°C for 14 hrs in a SW41Ti rotor (Beckman Coulter). After the gradients were fractioned into 24 to 26 0.5 ml aliquots, 20 µl of each fraction was analyzed by immunoblotting. If necessary, sucrose density gradient centrifugation was performed in the presence of GTPγS (20 mM) or GDP (20 mM).

**In vitro protein interaction assay.** IFT74, IFT81, and RABL2 cDNAs were amplified from total RNA of *Chlamydomonas* using the primer pairs (cIFT74-FOR and cIFT74-REV for IFT74, cIFT81-FOR and cIFT81-REV for IFT81, cRABL2-FOR and cRABL2-REV for RABL2) as listed in SI Appendix, Table S3. IFT81- and RABL2-expressing vectors were generated on pET28a (Novagen) backbones by double digestion with *Bam*HI and *Hin*dIII and DNA fragment ligation, resulting in pET28a-IFT81 and pET28a-RABL2. The desiring mutations Q83L and S37N in RABL2-encoding cDNA were introduced by site-directed mutagenesis using the primer pairs (cRABL2$^{Q83L}$-FOR and cRABL2$^{Q83L}$-REV for RABL2$^{Q83L}$ and cRABL2$^{S37N}$-FOR and cRABL2$^{S37N}$-REV for RABL2$^{S37N}$) as listed in SI Appendix, Table S3, resulting in pET28a-RABL2$^{Q83L}$ and pET28a-RABL2$^{S37N}$. These vectors were transformed into bacteria for expression of N-terminal 6×His-tagged recombinant proteins. IFT74-expressing vector was constructed on a pGEX-2T (GE Healthcare) to direct the expression of N-terminal GST-tagged IFT74 (GST-IFT74) (resulting in pGEX-2T-IFT74). The 6×His-tagged proteins and GST-IFT74 were purified with Ni-NTA and glutathione beads, respectively. The 6×His-tagged proteins were further cleaved with thrombin (Solarbio) to remove the N-terminal 6×His tag. To examine if RABL2 interacts with IFT74/81 directly, IFT81 and GST-IFT74 were mixed at 37°C for 2 hrs before undergoing glutathione sepharose purification for preparing the GST-IFT74/81 complex according to our previous report (9). GST-IFT74/81 or GST alone were next mixed with RABL2 in the presence of GTPγS (20 mM) or GDP (20 mM),

RABL2$^{Q83L}$, and RABL2$^{S37N}$. The mixture was incubated at 37°C for 2 hrs before glutathione sepharose purification was performed according to our previous report (9). Finally, 10 µg of eluted proteins were resolved on 12% SDS-PAGE gels followed by Coomassie staining for protein visualization. The generation of tag-free ARL3, ARL3$^{Q70L}$, and ARL3$^{T30N}$ recombinant proteins has been described in our previous report (6). The cDNAs encoding RABL2 and its mutants RABL2$^{Q83L}$ and RABL2$^{S37N}$ were cut from pET28a-RABL2, pET28a-RABL2$^{Q83L}$, pET28a-RABL2$^{S37N}$, respectively, and inserted into the *Bam*HI and *Hin*dIII sites of pGEX-2T (GE Healthcare), resulting vectors pGEX-2T-RABL2, pGEX-2T-RABL2$^{Q83L}$ and pGEX-2T-RABL2$^{S37N}$. These vectors were transformed into bacteria and were expressed as proteins with an N-terminal GST tag. ARL3, ARL3$^{Q70L}$, and ARL3$^{T30N}$ were mixed with the N-terminal GST tagged RABL2 (GST-RABL2) and its mutants (GST-RABL2$^{Q83L}$ and GST-RABL2$^{S37N}$) or GST alone, which were purified with glutathione sepharose beads. According to the experiments, glutathione sepharose purification was performed on the mixture as described previously (9). Ten micrograms of proteins from elutes were resolved on 12% SDS-PAGE gels and visualized with Coomassie staining. If necessary, the assay was carried out in the presence of GTPγS (20 mM), GDP (20 mM) or EDTA (20 µM).

**Size exclusion chromatography (SEC).** To determine if RABL2 and ARL3 bind each other and the molecule ratio of RABL2 to ARL3 in the transiently existing RABL2$^{GDP}$/ARL3$^{GDP}$ complex in vivo, ciliary matrix was prepared from 2.5 liters of CC-125 cells in the presence of GDP (20 mM) as described above and was equilibrated in 2M NaCl, HEPES (pH 7.0). The ciliary matrix solution was then fractionated on a HiLoad Superdex 16/600 200 pg column (GE Healthcare). To determine the molecule ratio of RABL2 to ARL3 in vitro, GST::RABL2$^{S37N}$/ARL3$^{T30N}$ complex obtained as described above was equilibrated in 2M NaCl, HEPES (pH 7.0) and fractionated on a HiLoad Superdex 16/600 200 pg column (GE Healthcare). For both in vitro and in vivo assays, a total of 180 fractions (1 ml/fraction) were collected. 20 µl of each of 13 fractions (65 to 77 ml for in vitro assays and 69 to 81 ml for in vivo assays) was analyzed by immunoblotting as described above. Protein standards (Blue Dextran (void volume), Aldolase (158 kD), Ovalbumin (44 kD), Ribonuclease A (13.7 kD), and Aprotinin (6.5 kD) were run on the same column equilibrated in 2M NaCl, HEPES (pH 7.0) to calculate the approximate size of complexes.

**In vitro GTP binding assay.** In vitro GTP binding assay was performed for determining GEF activity of RABL2 to ARL3 according to the previously described protocol (11). GTP affinities of bacterially expressed ARL3, RABL2, and mutants RABL2$^{Q83L}$ and RABL2$^{S37N}$ were determined by fluorescence spectrophotometric measurements of 2'(3')-O-(N-methylanthraniloyl)-labeled (mant-labeled) GTP (mant-GTP) (Abcam). To determine mant-GTP binding rate for GTPases, 50 µl of 2× binding reaction buffer (40 mM Tris pH 7.5, 100 mM NaCl, 20 mM MgCl2, and 1.5 µM mant-GTP), 4 µl of each GTPase or mutant alone (50 µM), 36 µl of pure water were added into each assigned well of a 96-well plate (black color to avid light) and mix well by gentle vortex. Time-dependent fluorescence change was recorded once every 30 sec (excitation at 355 nm and emission at 448 nm) using a SynergyH1/H1MF ELISA reader (BioTek). After 5 readings (150 seconds), 10 μl of each of the GEF candidates (5 μM) or pure water was added in wells, mixed well by pipetting, and continued to be read for another 1850 sec. Emission maxima of mant-GTP were plotted against time and normalized by nonlinear regression using GraphPad Prism 8.0 (GraphPad Software). Mant-GTP binding rate was calculated according to the formula: binding rate = Vmax (AFU/sec)/0.75 × Basal mant-GTP AFU × N. Vmax was determined by GraphPad Prism 8.0. Given that the N value for ARL3 is not available yet, we used the N value (N = 2) of CDC42 in all calculations as suggested previously (11).

**Living cell imaging and kymogram analysis.** Total internal reflection fluorescence (TIRF) microscopy was applied to visualize the motility of YFP-tagged IFT46, BBS8, RABL2, and its variants in cilia according to our previous report (7). The fluorescence proteins were imaged at ~15 frames per second (fps) using an IX83 inverted fluorescent microscopy (Olympus) equipped with a through-the-objective TIRF system, a 100×/1.49 NA TIRF oil immersion objective len (Olympus), a back illuminated scientific CMOS camera (Prime 95B, Photometrics), and 488-nm laser from Coherent OBIS Laser Module as detailed previously (7). Kymographs were generated and measured with CellSens Dimension (version 2.1, Olympus). IFT velocities and frequencies were calculated and presented using GraphPad Prism 8.0 (GraphPad Software).

**Measurement of ciliary length.** Phase contrast images of *Chlamydomonas* cell were taken by using an IX83 inverted fluorescent microscope (Olympus) equipped with a back illuminated scientific CMOS camera (Prime 95B, Photometrics) under a 60× objective. Ciliary length was measured according to our previous report (9).

**“In solution” mass spectrometry**

Protein complexes immunoprecipitated by camel anti-YFP antibody-conjugated agarose beads (V-nanoab Biotechnology) were visualized on Bolt Bis-Tris Plus Gels (NW04120BOX, Thermo Fisher Scientific) in Bolt MES SDS Running Buffer (B000202, Thermo Fisher Scientific) and visualized by silver staining as described previously (12). Mass spectrometry was performed by PTM Biolab. Inc. (Hangzhou, China). In details, the gel obtained above was cut into slices. The gel slices were destained in 50 mM $NH_4HCO_3$ (Sigma) in 50% acetonitrile (Fisher Commercial) (v/v) until clear. Gel pieces were dehydrated with 100 μl of acetonitrile for 5 min. After the liquid was removed, the gel pieces were incubated in 10 mM dithiothreitol (Sigma) at 56 °C for 60 min for rehydrating. The gel pieces were dehydrated again and rehydrated with 55 mM iodoacetamide (Sigma) by incubating at room temperature for 45 min in dark. Afterwards, the gel pieces were washed with 50 mM

$NH_4HCO_3$ and dehydrated with acetonitrile. The gel slices were added with 10 ng/µl trypsin (Promega) in 50 mM $NH_4HCO_3$ and sit on ice for 1 h followed by incubation at 37 °C overnight for in-gel digestion. After this, peptides were extracted with 50% acetonitrile, 5% formic acid (Fluka), and water followed by acetonitrile, freeze-dried, and reconstituted in 2% acetonitrile, 0.1% formic acid, and water. Prior to LC-MS, EASY-nLC 1000 UPLC (ThermoFisher Scientific) LC instrument was set at a flow rate of either 550 nL/min where mobile phase A was 0.1% formic acid in water and mobile phase B was 0.1% formic acid in 98% acetonitrile. Peptide pools were directly injected onto the home-made reversed-phase analytical column in which linear gradients (6-35% B, 35-80% B, and constant 80%) were of 22 min, 4 min, and 4 min eluting peptides into the mass spectrometer. The Q Exactive™ Plus (ThermoFisher Scientific) mass spectrometer were used, where a ''fastest'' MS/MS data acquisition was used. MS/MS was acquired using DDA with a collisional energy of 28. Raw files were processed using Proteome Discoverer 2.1 (ThermoFisher Scientific) using 10 ppm mass accuracy limits for precursors and 0.02 Da mass accuracy limits for MS/MS spectra. MS/MS data was compared to the *Chlamydomonas* Flagellar Proteome Data (http://chlamyfp.org/readcsvfile_js_Pfam.php). Spectral counts allow up to two missed cleavages per peptide. All data was filtered and presented at a 1% false discovery rate. Post processing using Excel was used to further process the data for gene ontological analyses.

**Population phototaxis assay.** Population phototaxis assays were performed on *Chlamydomonas* cells according to the protocol reported previously (13). In brief, *Chlamydomonas* cells growing to a concentration of ~$10^7$ cells were harvested and 100 µl of the cell suspension were placed into the surface of Petri dishes of 3.5-cm diameter (706001; Wuxi NEST Biotech.) containing solid TAP medium. Afterwards, the cells were illuminated with a flashlight from one side for 4 min. Images were continuously taken once every two minutes with a standard digital camera (Nikon A70).

**Single-cell motion assay.** Single-cell motion assay was performed on *Chlamydomonas* cells according to the protocol reported previously (13). Briefly, 20 µl of cell suspensions as obtained above were placed on superfrost™ plus microscope slides (12-550-15; Fisher Brand) and observed using an inverted microscope (IX83, Olympus) under non-phototactic red light illumination (561-nm laser from Coherent OBIS Laser Module). The cells were then illuminated for 5 sec with phototactic active flashlight. Afterwards, five images were continuously taken once every 0.3 sec using a back illuminated scientific CMOS camera (Prime 95B, Photometrics). The five sequential images each displayed in a different color were merged in ImageJ (version 1.42g, National Institutes of Health) to show the swimming tracks of single cells, allowing us to determine the angle and the direction of a cell's movements. Excel for Mac (version 16.52) were applied to analyze the data and generate polar histograms with 60° bins.

**Statistics.** Statistical analysis was performed with GraphPad Prism 8.0 (GraphPad Software). If not otherwise specified, all data were shown as mean ± S.D. from three independently performed experiments and n means sample numbers. For comparisons on velocities and frequencies of the YFP-tagged proteins, one-sample unpaired student *t*-test was used. For comparisons on expression levels of protein and the ciliary length, two-tailed *t*-test was used. If not mentioned elsewhere, significance was set as $p < 0.001$ (marked as **). "n.s." represents non-significance.

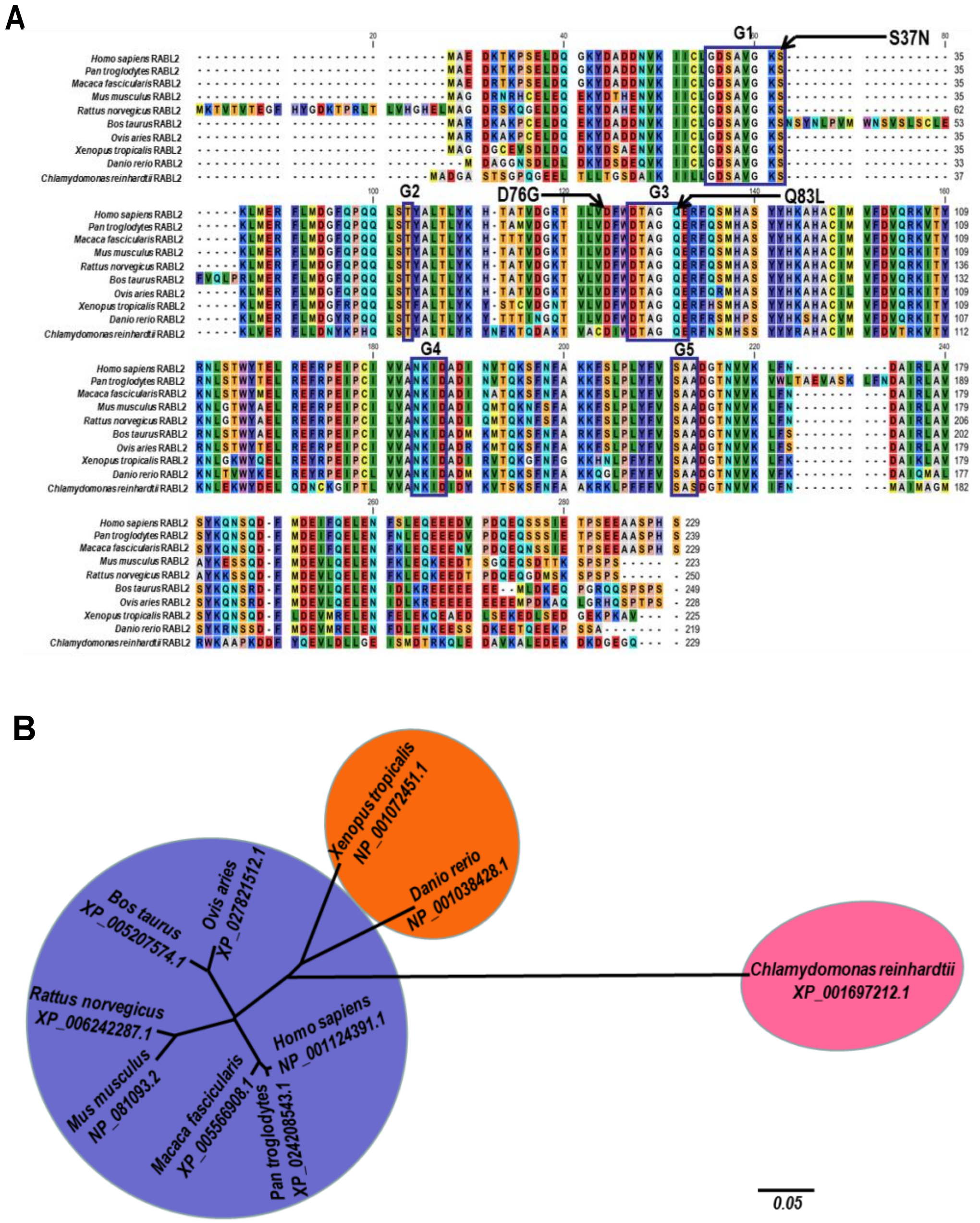


**Fig. S1.** RABL2 is highly conserved across ciliated species. (*A*). Sequence alignment of deduced amino acid sequences from ten invertebrate and vertebrate RABL2 orthologues. Alignments were generated using CLC main workbench (version 6.8); the most conserved residues are shown in black, the least conserved are in red. Dashes indicate gaps introduced to optimize the alignment. (B). The phylogenetic tree of RABL2 proteins from invertebrate and vertebrate species as indicated. The neighbor-joining tree was calculated using the TreeView 1.6.6 software. Branch length represents evolutionary relatedness. Accession numbers are as follows: *Bos taurus*, XP_005207574.1; *Ovis aries*, XP_027821512.1; *Homo sapiens*, NP_001124391.1; *Xenopus tropicalis*, NP_001072451.1; *Rattus norvegicus*, XP_006242287.1; *Danio rerio*, NP_001038428.1;

*Mus musculus*, NP_081093.2; *Macaca fascicularis*, XP_005566908.1; *Pan Troglodytes*, XP_024208543.1; and *Chlamydomonas reinhardtii*, XP_001697212.1.

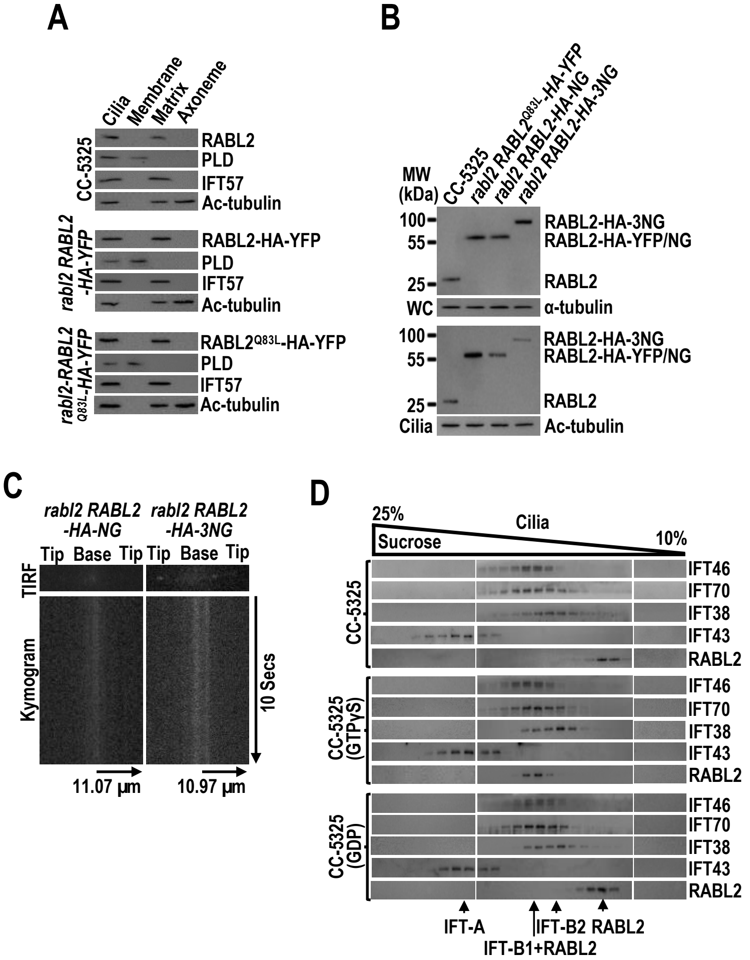


**Fig. S2.** RABL2$^{GTP}$ binds IFT-B1 in cilia (related to Fig. 3). (*A*). Immunoblots of ciliary fractions prepared from cells indicated on the left probed with α-RABL2, α-PLD (ciliary membrane marker), α-IFT57 (ciliary matrix marker), and acetylated (Ac)-tubulin (axoneme marker). (*B*). Immunoblots of whole cell (WC) and ciliary samples prepared from cells indicated on the top probed with α-RABL2. Alpha-tubulin and acetylated (Ac)-tubulin were used for adjusting the loading for WC samples and cilia, respectively. MW stands for molecular weight. (*C*). TIRF images and corresponding kymograms of cells indicated on the top (Movies S4-S5, 15 fps). The time and transport lengths are indicated on the right and on the bottom, respectively. The ciliary base (base) and tip (tip) were shown. (*D*). Immunoblots of sucrose density gradients of CC-5325 cilia in the presence of GTPγS, GDP, or none of them probed for the IFT-B1 subunits IFT46 and IFT70, the IFT-B2 subunit IFT38, the IFT-A subunit IFT43, and RABL2. RABL2 co-sediments with IFT-B1 only in the presence of GTPγS.

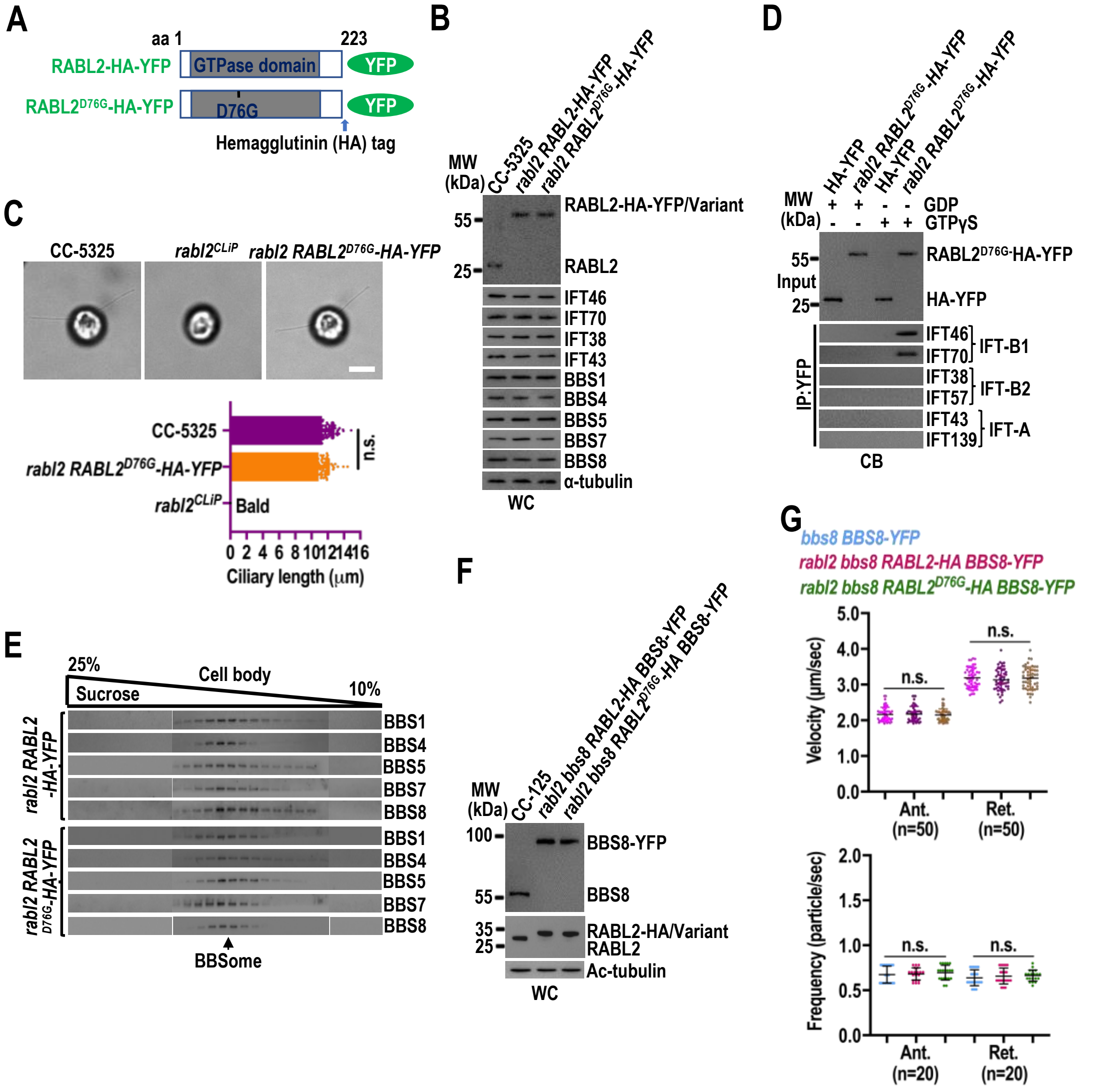


**Fig. S3.** RABL2$^{D76G}$ rescues ciliation but fails to enter cilia for promoting BBSome movement out of cilia (related to Fig. 4). (*A*) Schematic representation of RABL2$^{D76Q}$-HA-YFP as compared to RABL2-HA-YFP. aa stands for amino acid position. YFP: yellow fluorescence protein. The hemagglutinin (HA) tag was shown. (*B*) Immunoblots of WC samples prepared from cells indicated on the top probed with α-RABL2, α-IFT46, α-IFT70, α-IFT38, α-IFT43, α-BBS1, α-BBS4, α-BBS5, α-BBS7, and α-BBS8. (*C*) Representative phase contrast (PC) images (Upper) of CC-5325, *rabl2$^{CLiP}$*, and *rabl2 RABL2$^{D76G}$-HA-YFP* cells were shown. Scale bars: 5 µm. (Bottom) *rabl2* RABL2$^{D76G}$-HA-YFP (10.16 ± 1.11 µm, n = 100) cells had full-length cilia as compared to CC-5325 (10.31 ± 0.75 µm, n = 100) cells (right). *rabl2$^{CLiP}$* cells were bald. Mean lengths are listed; error bar indicates S.D. and "n" indicates the number of cilia counted. (*D*) Immunoblots of α-YFP-captured proteins from cell body (CB) samples prepared from cells indicated on the top in the presence of GTPγS or GDP probed for the IFT-B1 subunits IFT46 and IFT70, the IFT-B2 subunits IFT38 and IFT57, the IFT-A subunits IFT43 and IFT139. Input was quantified with α-YFP by immunoblotting.

(*E*) Immunoblots of sucrose density gradients of CB samples prepared from cells indicated on the left probed for the BBSome subunits BBS1, BBS4, BBS5, BBS7, and BBS8. (*F*) Immunoblots of WC samples prepared from cells indicated on the top probed with α-BBS8 and α-RABL2. (*G*) Velocities and frequencies of BBS8-YFP to traffic inside cilia of cells indicated on the top shown as graphs. Mean values are listed. Error bar indicates S.D. n: number of cilia analyzed. n.s.: non-significance. For panels *B* and *F*, α-tubulin was used as a loading control. For panels *B*, *D*, and *F*, MW stands for molecular weight.

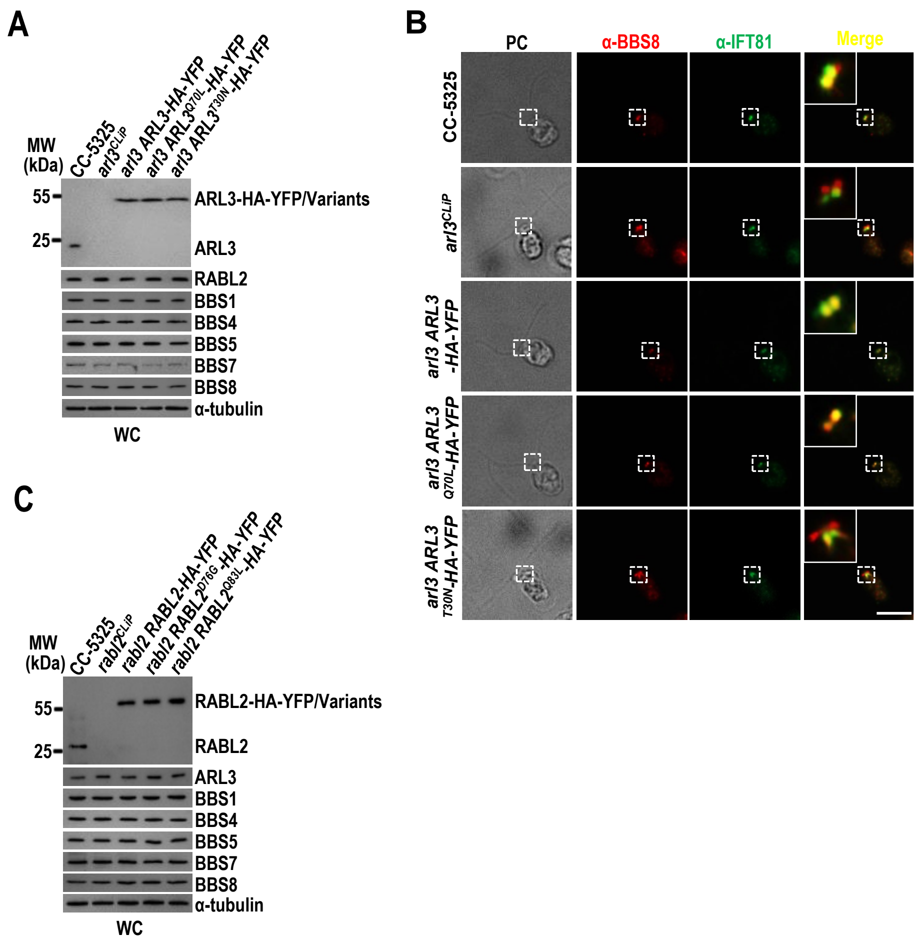


**Fig. S4**. RABL2 mediates outward BBSome movement cross the TZ by activating ARL3 (related to Fig. 5). (*A*) Immunoblots of WC samples prepared from cells indicated on the top probed with α-ARL3, α-RABL2, α-BBS1, α-BBS4, α-BBS5, α-BBS7, and α-BBS8. (*B*) Cell indicated on the left stained with α-BBS8 (red) and α-IFT81 (green). PC images of cells were shown. Inset shows the basal bodies and the transition zone staining. Scale bars: 10 µm. (*C*) Immunoblots of WC samples prepared from cells indicated on the top probed with α-RABL2, α-ARL3, α-BBS1, α-BBS4, α-BBS5, α-BBS7, and α-BBS8. For panels *A* and *C*, α-tubulin was used as a loading control. MW stands for molecular weight.

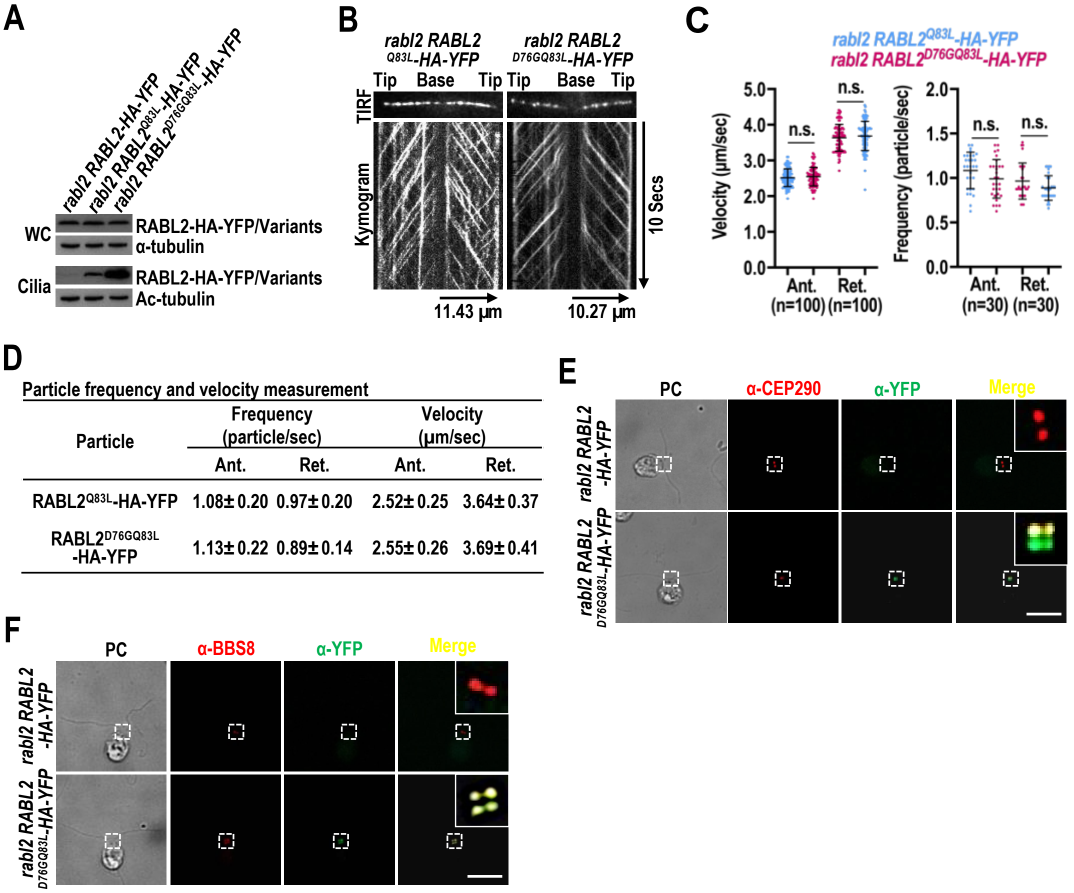


Particle frequency and velocity measurement

| Particle | Frequency (particle/sec) Ant. | Frequency (particle/sec) Ret. | Velocity (μm/sec) Ant. | Velocity (μm/sec) Ret. |
|---|---|---|---|---|
| RABL2$^{Q83L}$-HA-YFP | 1.08±0.20 | 0.97±0.20 | 2.52±0.25 | 3.64±0.37 |
| RABL2$^{D76GQ83L}$-HA-YFP | 1.13±0.22 | 0.89±0.14 | 2.55±0.26 | 3.69±0.41 |

**Fig. S5.** RABL2$^{D76GQ83L}$ undergoes normal IFT and retains the ability to shed from retrograde IFT at the TZ region but fails to move cross the TZ for ciliary retrieval. (*A*) Immunoblots of WC and ciliary samples prepared from cells indicated on the top probed with α-RABL2. Alpha-tubulin and acetylated (Ac)-tubulin were used for adjusting the loading for WC samples and cilia, respectively. MW stands for molecular weight. (*B*) TIRF images and corresponding kymograms of cells indicated on the top (Movies S9 and S10, 15 fps). The time and transport lengths are indicated on the right and on the bottom, respectively. The ciliary base (base) and tip (tip) were shown. (*C* and *D*) Velocities and frequencies of YFP-tagged proteins to traffic inside cilia of cells indicated on the top shown as graphs (*C*) and numbers (*D*). Mean values are listed. Error bar indicates S.D. from 100 and 30 cilia analyzed for determining velocity and frequency, respectively. (*E*) Cells indicated on the left stained with α-CEP290 (red) and α-IFT81 (green). (*F*) Cells indicated on the left stained with α-BBS8 (red) and α-YFP (green). For panels *E* and *F*, PC images of cells were shown. Inset shows the basal bodies and the transition zone staining. Scale bars: 10 μm.

**Table S1**. Antibodies used in this study.

| Antibody | Dilution | | Origins | Reference or source |
|---|---|---|---|---|
| | IB | IS | | |
| Anti-RABL2 | 1:500 | N/A | Rabbit | This study |
| Anti-ARL3 | 1:250 | N/A | Rabbit | (6) |
| Anti-CEP290 | N/A | 1:100 | Rabbit | (6) |
| Anti-IFT38 | 1:1,000 | 1:200 | Rabbit | (10) |
| Anti-IFT46 | 1:1,000 | 1:200 | Rabbit | (5) |
| Anti-IFT57 | 1:500 | N/A | Rabbit | (5) |
| Anti-IFT43 | 1:500 | 1:200 | Rabbit | (14) |
| Anti-IFT70 | 1:1,000 | 1:200 | Rabbit | (5) |
| Anti-IFT81 | N/A | 1:200 | Mouse | (9) |
| Anti-IFT139 | 1:1,000 | N/A | Rabbit | (5) |
| Anti-BBS1 | 1:1,000 | N/A | Rabbit | (7) |
| Anti-BBS4 | 1:500 | N/A | Rabbit | (2) |
| Anti-BBS5 | 1:1,000 | N/A | Rabbit | (7) |
| Anti-BBS7 | 1:500 | N/A | Rabbit | (2) |
| Anti-BBS8 | 1:500 | 1:100 | Rabbit | (10) |
| Anti-PLD | 1:1,000 | N/A | Rabbit | (2) |
| Anti-α-tubulin | 1:10,000 | N/A | Mouse | Sigma-Aldrich |
| Anti-acetylated-tubulin | 1:30,000 | N/A | Mouse | Sigma-Aldrich |
| Anti-GFP (YFP) | 1:1,000 | 1:50 | Mouse | Roche |
| HRP-conjugated goat anti-rabbit IgG | 1:10,000 | N/A | Goat | The Jackson Lab. |
| HRP-conjugated goat anti-mouse IgG | 1:10,000 | N/A | Goat | The Jackson Lab. |
| HRP-conjugated goat anti-rat IgG | 1:10,000 | N/A | Goat | The Jackson Lab. |
| Alexa-Fluor 594-conjugated goat anti-rabbit IgG | N/A | 1:400 | Goat | Molecular Probes |
| Alexa-Flour 488-conjugated goat anti-mouse IgG | N/A | 1:400 | Goat | Molecular Probes |
| Alexa-Fluor 594-conjugated goat anti-mouse IgG | N/A | 1:400 | Goat | Molecular Probes |

**Table S2.** *Chlamydomonas* strains used in this study

| Name | Genotype | Reference or source |
|---|---|---|
| CC-125 (wild-type) | *nit1; nit2; mt*$^{+}$ | CRC |
| CC-5325 | *cw15; mt*$^{-}$ | CLiP (15) |
| *rabl2*$^{CLiP}$ | *cw15*; *rabl2*; *mt*$^{-}$ | CLiP (15) |
| *arl3*$^{CLiP}$ | *cw15*; *arl3*; *mt*$^{-}$ | CLiP (15) |
| *bbs8* | *bbs8*; *mt*$^{+}$ | (16) |
| *bbs8 BBS8-YFP* | *bbs8*; *mt*$^{+}$*; BBS8-YFP-TG* | (6) |
| *rabl2 RABL2-HA-YFP* | *cw15*; *rabl2*; *mt*$^{-}$*; RABL2-HA-YFP-TG* | This study |
| *rabl2 RABL2*$^{Q83L}$*-HA-YFP* | *cw15*; *rabl2*; *mt*$^{-}$*; RABL2*$^{Q83L}$*-HA-YFP-TG* | This study |
| *rabl2 RABL2*$^{S37N}$*-HA-YFP* | *cw15*; *rabl2*; *mt*$^{-}$*; RABL2*$^{S37N}$*-HA-YFP-TG* | This study |
| *rabl2 RABL2*$^{D76G}$*-HA-YFP* | *cw15*; *rabl2*; *mt*$^{-}$*; RABL2*$^{D76G}$*-HA-YFP-TG* | This study |
| *rabl2 RABL2*$^{D76GQ83L}$*-HA-YFP* | *cw15*; *rabl2*; *mt*$^{-}$*; RABL2*$^{D76GQ83L}$*-HA-YFP-TG* | This study |
| *rabl2 RABL2-HA-NG* | *cw15*; *rabl2*; *mt*$^{-}$*; RABL2-HA-NG-TG* | This study |
| *rabl2 RABL2-HA-3NG* | *cw15*; *rabl2*; *mt*$^{-}$*; RABL2*$^{L}$*-HA-3NG-TG* | This study |
| *ift46-1 IFT46-YFP* | *cw15*; *rabl2*; *mt*$^{-}$*; IFT46-YFP-TG* | (17) |
| *arl3 ARL3-HA-YFP* | *cw15*; *arl3*; *mt*$^{-}$*; ARL3-HA-YFP-TG* | (6) |
| *arl3 ARL3*$^{Q70L}$*-HA-YFP* | *cw15*; *arl3*; *mt*$^{-}$*; ARL3*$^{Q70L}$*-HA-YFP-TG* | (6) |
| *arl3 ARL3*$^{T30N}$*-HA-YFP* | *cw15*; *arl3*; *mt*$^{-}$*; ARL3*$^{t30N}$*-HA-YFP-TG* | (6) |
| *rabl2 RABL2*$^{Q83L}$*-HA-YFP ARL3*$^{Q70L}$*-HA-YFP* | *cw15*; *rabl2*; *mt*$^{-}$*; RABL2*$^{Q83L}$*-HA-YFP-TG; ARL3*$^{Q70L}$*-HA-YFP-TG* | This study |
| *rabl2 RABL2-HA* | *cw15*; *rabl2*; *mt*$^{-}$*; RABL2-HA-TG* | This study |
| *rabl2 RABL2*$^{Q83L}$*-HA* | *cw15*; *rabl2*; *mt*$^{-}$*; RABL2*$^{Q83L}$*-HA-TG* | This study |
| *rabl2 RABL2*$^{D76G}$*-HA* | *cw15*; *rabl2*; *mt*$^{-}$*; RABL2*$^{D76G}$*-HA-TG* | This study |
| *rabl2 RABL2*$^{D76GQ83L}$*-HA* | *cw15*; *rabl2*; *mt*$^{-}$*; RABL2*$^{D76GQ83L}$*-HA-TG* | This study |
| *rabl2 bbs8 RABL2-HA BBS8-YFP* | *rabl2*; *bbs8*; *RABL2-HA-TG; BBS8-YFP-TG* | This study |
| *rabl2 bbs8 RABL2*$^{D76G}$*-HA BBS8-YFP* | *rabl2*; *bbs8*; *RABL2*$^{D76G}$*-HA-TG; BBS8-YFP-TG* | This study |
| *rabl2 bbs8 RABL2*$^{Q83L}$*-HA BBS8-YFP* | *rabl2*; *bbs8*; *RABL2*$^{Q83L}$*-HA-TG; BBS8-YFP-TG* | This study |
| *rabl2 bbs8 RABL2*$^{D76GQ83L}$*-HA BBS8-YFP* | *rabl2*; *bbs8*; *RABL2*$^{D76GQ83L}$*-HA-TG; BBS8-YFP-TG* | This study |
| *rabl2 bbs8 RABL2-HA-YFP* | *rabl2*; *bbs8*; *RABL2-HA-YFP-TG* | This study |
| *rabl2 bbs8 RABL2*$^{D76G}$*-HA-YFP* | *rabl2*; *bbs8*; *RABL2*$^{D76G}$*-HA-YFP-TG* | This study |
| *rabl2 bbs8 RABL2*$^{Q83L}$*-HA-YFP* | *rabl2*; *bbs8*; *RABL2*$^{Q83L}$*-HA-YFP-TG* | This study |
| *arl3 bbs8 ARL3-HA-YFP* | *arl3*; *bbs8*; *ARL3-HA-YFP-TG* | (6) |
| *arl3 bbs8 ARL3*$^{Q70L}$*-HA-YFP* | *arl3*; *bbs8*; *ARL3*$^{Q70L}$*-HA-YFP-TG* | (6) |
| *arl3 bbs8 ARL3*$^{T30N}$*-HA-YFP* | *arl3*; *bbs8*; *RL3*$^{T30N}$*-HA-YFP-TG* | (6) |
| HR-YFP | *nit1; nit2; mt*$^{+}$*; HA-YFP-TG* | (18) |
| RABL2$^{miRNA}$ | *nit1; nit2; mt*$^{+}$*; RABL2*$^{miRNA}$ | This study |
| RABL2$^{Res\text{-}WT}$ | *nit1; nit2; mt*$^{+}$*; RABL2*$^{miRNA}$*; RABL2-HA-YFP-TG* | This study |
| RABL2$^{Res\text{-}D76G}$ | *nit1; nit2; mt*$^{+}$*; ARL3*$^{miRNA}$*; RABL2*$^{Q83L}$*-HA-YFP-TG* | This study |
| RABL2$^{Res\text{-}Q83L}$ | *nit1; nit2; mt*$^{+}$*; ARL3*$^{miRNA}$*; RABL2*$^{S37N}$*-HA-YFP-TG* | This study |

Note: CRC stands for *Chlamydomonas* Resource Center.

**Table S3.** Primers used in this study

| Name | Nucleotide sequence |
|---|---|
| **Primers used to clone target genes** | |
| gRABL2-FOR | 5'-GCTCTAGACAACGCAGCTCACTGGAG-3' |
| gRABL2-REV | 5'-CGGAATTCTTGCCCCTCGCCGTCC-3' |
| gRABL2::HA-REV | 5'-CCGCTCGAGTTAGGCGTAGTCGGGCACGTCGTAGGGGTATTGCCCCTCGCCGTCC-3' |
| HYG-FOR | 5'-GTATTGAAGCGGGGCCCGGTACCCTTTCTTGCGCTATGACAC-3' |
| HYG-REV | 5'-CTCACTATAGGGCGAATTGGGTACCCCATGGGATGACGGGCCCGG-3' |
| gARL3$^{Q70L}$-FOR | 5'-CCGCGGTGGCGGCCGCTCTAGACCGACATGAGCGCCTC-3' |
| gARL3$^{Q70L}$-REV | 5'-GGGGGATCCACTAGTTCTAGACTTGACCTGC TTCATC-3' |
| cRABL2-FOR | 5'-CGGGATCCATGGCTGATGGGGCGAGC-3' |
| cRABL2-REV | 5'-CCCAAGCTTTTATTGCCCCTCGCCGTCC-3' |
| HA::NG-FOR | 5'-GGAATTCTACCCCTACGACGTGCCCGACTACGCCATGGTGAGCAAGGGCGAG-3' |
| HA::NG-REV | 5'-CCGCTCGAGTTACTTGTACAGCTCGTCC-3' |
| NG-1-FOR | 5'-GGAATTCTACCCCTACGACGTGCCCGACTACGCCATGGTGAGCAAGGGCGAG-3 |
| NG-1-REV | 5'-CGCGGATCCCTTGTACAGCTCGTCCATGCC-3' |
| NG-2-FOR | 5'-CGCGGATCCATGGTGAGCAAGGGCGAG-3' |
| NG-2-REV | 5'-CCCAAGCTTCTTGTACAGCTCGTCCATGCC-3' |
| NG-3-FOR | 5'-CCCAAGCTTATGGTGAGCAAGGGCGAGGAG-3' |
| NG-3-REV | 5'-CCGCTCGAGTTACTTGTACAGCTCGTCC-3' |
| cIFT74-FOR | 5'-CGCGGATCCATGGACAGGCCCTCTAGCCGCGG-3' |
| cIFT74-REV | 5'-CCCAAGCTTTTACACCACGGCCTTCTTGAC-3' |
| cIFT81-FOR | 5'-CGCGGATCCATGGGAGACGTGAGCTACATCG-3' |
| cIFT81-REV | 5'-CCCAAGCTTTTAGAGCTGCAGCACGTTGGCGC-3' |
| **Primers used to do site-directed mutagenesis** | |
| RABL2$^{Q83L}$-FOR | 5'-GACACGGCCGGGCTGGAGCGCTTCAAC-3' |
| RABL2$^{Q83L}$-REV | 5'-CTGTGCCGGCCCGACCTCGCGAAGTTG-3' |
| RABL2$^{S37N}$-FOR | 5'-CTCCGCTGTGGGGAAAAACAAGTAAGCTCAGCGGTC-3' |
| RABL2$^{S37N}$-REV | 5'-GAGGCGACACCCCTTTTTGTTCATTCGAGTCGCCAG-3' |
| RABL2$^{D76G}$-FOR | 5'-GCTGCTCGGACAGGCATATGGGAC-3' |
| RABL2$^{D76G}$-REV | 5'-CGACGAGCCTGTCCGTATACCCTG-3' |
| cRABL2$^{Q83L}$-FOR | 5'-GACACGGCCGGGCTGGAGCGCTTCAAC-3' |
| cRABL2$^{Q83L}$-REV | 5'-CTGTGCCGGCCCGACCTCGCGAAGTTG-3' |
| cRABL2$^{S37N}$-FOR | 5'-CTCCGCTGTGGGGAAAAACAAGCTCGTGG-3' |
| cRABL2$^{S37N}$-REV | 5'-GAGGCGACACCCCTTTTTGTTCGAGCACC-3' |
| **Primers used to generate RABL2 miRNA construct** | |
| RABL2 miRNA-3'-UTR | 5'-GATATCAGGAAACCAAGGCGCGCTAGCTTCCTGGGCGCAGTGTTCCAGCT<br>GCAGTACTTATGACGCAAACGTCGACACCTCGCTGATCGGCACCATGGGG<br>GTGGTGGTGATCAGCGCTAGTGTCGACGTTTGCGTCATAATACTGCAGCC<br>GGAACACTGCCAGGAGAATTC-3' |

**Movie S1.** TIRF imaging of IFT46-YFP movement in *ift46-1 IFT46-YFP* cilia. A frame from this movie and kymograph are shown in Figure 3*B*. Play speed is real-time (15 fps).

**Movie S2.** TIRF imaging of RABL2$^{Q83L}$-HA-YFP movement in *rabl2 RABL2$^{Q83L}$-HA-YFP* cilia. A frame from this movie and kymograph are shown in Figure 3*B*. Play speed is real-time (15 fps).

**Movie S3.** TIRF imaging of RABL2-HA-YFP movement in *rabl2 RABL2-HA-YFP* cilia. A frame from this movie and kymograph are shown in Figure 3*B*. Play speed is real-time (15 fps).

**Movie S4.** TIRF imaging of RABL2-HA-NG movement in *rabl2* RABL2-HA-NG cilia. A frame from this movie and kymograph are shown in Figure S2*C*. Play speed is real-time (15 fps).

**Movie S5.** TIRF imaging of RABL2-HA-3NG movement in *rabl2* RABL2-HA-3NG cilia. A frame from this movie and kymograph are shown in Figure S2*C*. Play speed is real-time (15 fps).

**Movie S6.** TIRF imaging of BBS8-YFP movement in *bbs8* BBS8-YFP cilia. A frame from this movie and kymograph are shown in Figure 4*E*. Play speed is real-time (15 fps).

**Movie S7.** TIRF imaging of BBS8-YFP movement in *rabl2 bbs8 RABL2-HA BBS8-YFP* cilia. A frame from this movie and kymograph are shown in Figure 4*E*. Play speed is real-time (15 fps).

**Movie S8.** TIRF imaging of BBS8-YFP movement in *rabl2 bbs8 RABL2$^{D76G}$-HA BBS8-YFP* cilia. A frame from this movie and kymograph are shown in Figure 4*E*. Play speed is real-time (15 fps).

**Movie S9.** TIRF imaging of RABL2$^{Q83L}$-HA-YFP movement in *rabl2 RABL2$^{Q83L}$-HA-YFP* cilia. A frame from this movie and kymograph are shown in Figure S5*B*. Play speed is real-time (15 fps).

**Movie S10.** TIRF imaging of RABL2$^{D76GQ83L}$-HA-YFP movement in *rabl2 RABL2$^{D76GQ83L}$-HA-YFP* cilia. A frame from this movie and kymograph are shown in Figure S5*B*. Play speed is real-time (15 fps).
